# Signatures of kin selection in a natural population of the bacteria *Bacillus subtilis*

**DOI:** 10.1101/2022.11.07.515416

**Authors:** Laurence J. Belcher, Anna E. Dewar, Chunhui Hao, Melanie Ghoul, Stuart A. West

## Abstract

Laboratory experiments have suggested that bacteria perform a range of cooperative behaviours, which are favoured because they are directed towards relatives (kin selection). However, there is a lack of evidence for cooperation and kin selection in natural bacterial populations. Molecular population genetics offers a promising method to study natural populations, because theory predicts that kin selection will lead to relaxed selection, which will result in increased polymorphism and divergence at cooperative genes. Examining a natural population of *Bacillus subtilis*, we found consistent evidence that putatively cooperative traits have higher polymorphism and greater divergence than putatively private traits expressed at the same rate. In addition, we were able to eliminate alternative explanations for these patterns, and found more deleterious mutations in genes controlling putatively cooperative traits. Overall, our results suggest cooperation favoured by kin selection, with an average relatedness of *r*=0.77 between interacting individuals.

## Introduction

Laboratory studies have suggested that bacteria cooperate in a diversity of ways (1, 2). Individual cells produce and secrete molecules to collectively scavenge nutrients, fight antibiotics, and move through their environment (3–5). This cooperation is however vulnerable to cheating by non-producers, who withhold their own cooperation whilst benefiting from that of others (6). A resolution to this vulnerability is kin selection, where cooperation is favoured because the benefits of cooperation go to related cells which share the gene for cooperation (7). Laboratory experiments have also supported a role of kin selection, with experimental evolution, and by showing how clonal growth makes neighbouring cells highly related, and limited diffusion keeps secreted molecules in the neighbourhood (3, 8–11).

In contrast, there is little evidence for cooperation and kin selection in natural populations of bacteria (12–15). The extent to which bacteria cooperate and interact with close relatives is likely to be highly dependent upon environmental conditions. It is hard to know whether the artificial environments and gene knockouts of lab experiments are representative of natural populations. Experiments have also shown that some traits can be cooperative in some environments but private in others (16). Across species comparative studies have shown that cooperation is more common in species where relatedness is higher (17, 18), but this doesn’t help us determine whether specific traits are evolving as cooperative public goods. For this, we need a way to study bacteria in their natural environment.

Molecular population genetics offers a promising method to test for cooperation and kin selection in natural populations. Theory predicts that kin selection leaves a distinct signature (footprint) of selection in genomes (Figure 1) (19–23). If a gene encodes a trait that provides a direct benefit to the individual performing that trait, then the cells which perform that trait carry the gene for that trait. Similarly, in a clonal population, relatedness between cells is *r* = 1, and so the cells which benefit from cooperation also carry the cooperative gene. If instead, relatedness between cells is *r* < 1, the cells which benefit from cooperation won’t always carry the gene for cooperation. This dilution of the benefit of cooperation leads to relaxed (reduced) selection, relative to directly beneficial traits or in a clonal population (*r* = 1).

**Figure 1:**
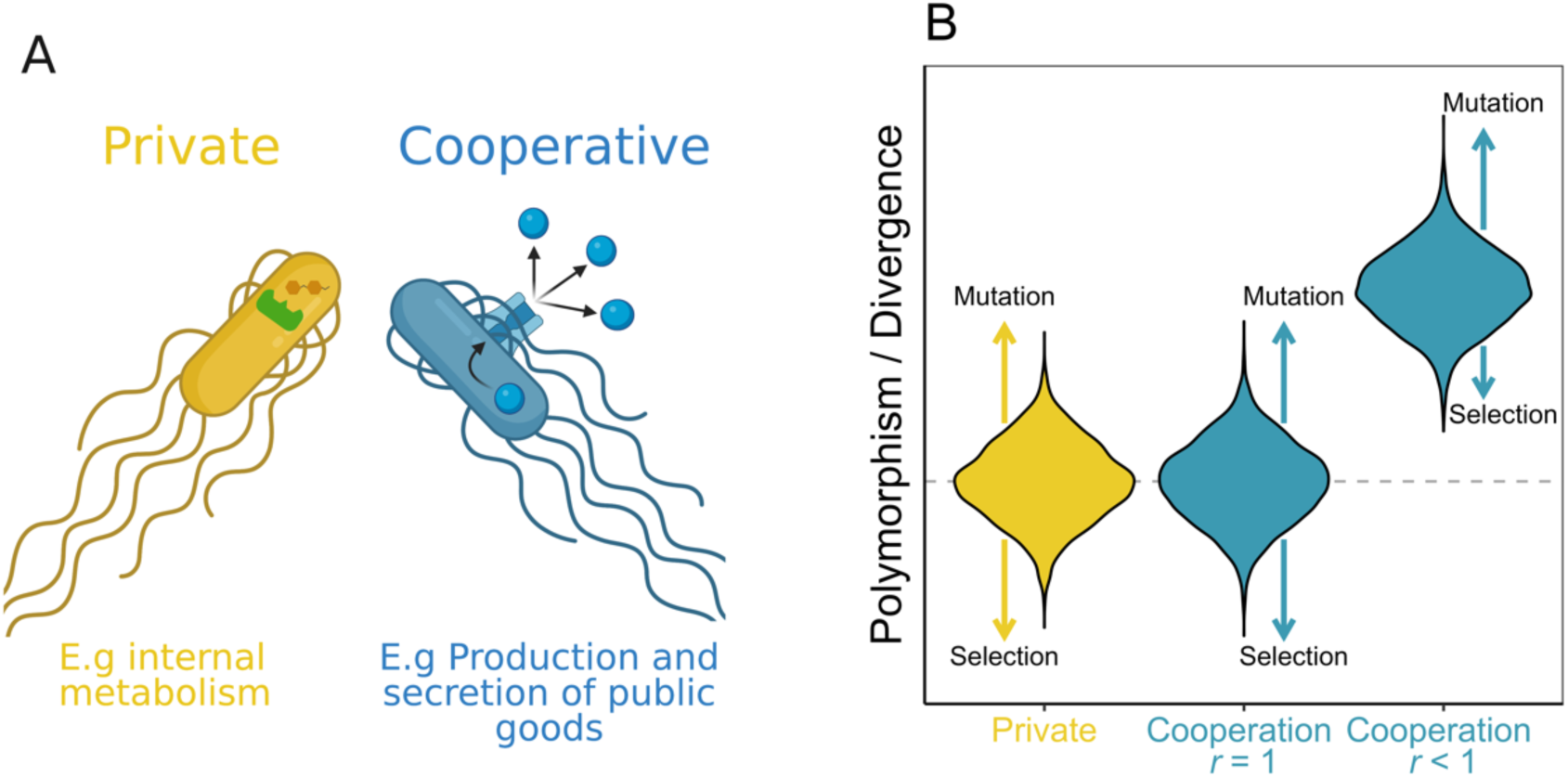
Population genetic theory for cooperative traits. (A) Representation of the categorisation of traits as private (yellow) or cooperative (blue). Cooperative traits involve the production and secretion of molecules whose fitness benefits are shared with other nearby cells. Private traits are those whose fitness benefits are only felt by the individuals expressing the trait (B) Prediction for relative polymorphism and divergence for private (yellow) and cooperative (blue) genes. If relatedness, *r*=1, then cooperative genes (middle blue violin) should have the same polymorphism and divergence as private genes (left yellow violin). In contrast, if *r*<1 then cooperative genes (right blue violin) should show greater polymorphism and divergence than private genes. Figure based on (19, 21).

The relaxed selection when *r* < 1 results in an increased probability of fixation for deleterious mutations, and a decreased probability of fixation for beneficial mutations (14–16). The consequence of this change in fixation probabilities, when *r*<1, is that it would lead to increased polymorphism and divergence in cooperative genes relative to genes that have direct fitness effects (Figure 1). Consequently, by examining patterns of polymorphism and divergence, we can test for signatures of cooperation favoured by kin selection. This method has been applied in microbes to the social amoeba *Dictyostelium discoideum* with mixed results, although relatedness is close to *r* = 1 in that species, and so we might not expect to a significant signature of kin selection (24–27). We have also previously applied this method to the opportunistic pathogen *Pseudomonas aeruginosa*, and found evidence for cooperative traits favoured by kin selection (28). A key advantage of this method is that it solves the problem caused by traits that are only cooperative in certain contexts, as the signature of selection will represent the average conditions experienced over time.

However, there were two potential problems with the only previous analysis with bacteria, on *P. aeruginosa* (28). Firstly, the genomes had been collected over several decades, and from six different continents. The underlying population genetic theory assumes that all genomes are sampled from a single population, at a single time point. Violating this assumption could have led to biased or spurious results (29). Secondly, we were unable to directly control for genes having conditional fitness effects. If a gene only has fitness effects in certain environments or certain generations, then this can also lead to relaxed selection, with increased polymorphism and divergence (21). We can control for this by looking at gene expression data to see if the genes we are comparing tend to be expressed together. Belcher *et al*. lacked data on gene expression, and so controlled for this problem by making targeted comparisons between cooperative and private traits that they argued were likely to be expressed at different rates (28). If their assumptions did not hold, and expression rates differed, then then their patterns could alternatively be explained by gene expression rather than kin selection for cooperation.

We were able to address these problems by taking advantage of two recently developed data sets in the bacteria *Bacillus subtilis*. *B. subtilis* is found in soil and the gastrointestinal tracts of several animals, including humans, and is used on an industrial scale by biotechnology companies (30). A number of laboratory studies have suggested that *B. subtilis* is a highly cooperative species, which secretes a number of potentially cooperative enzymes (31–34). First, we used a natural population consisting of 31 environmental isolates collected as part of a citizen science project in Dundee, Scotland (35) (Supplementary Table 1). All of these strains were collected from the wild around the same time, and from similar niches. Second, we were able to directly control for gene expression rates, by using two genome-wide studies of gene expression across several timepoints during biofilm formation (36, 37). Taken together, these studies allow us to satisfy the assumptions of the underlying population genetic theory.

## Results & Discussion

We compared genes controlling traits that are hypothesised to be cooperative, with traits that are hypothesised to be private. We identified six different types of putatively cooperative behaviour, where an appropriate comparison could be made with genes controlling private traits, that are likely to be expressed at similar rates (Table 1). We used the first type, quorum sensing (QS), for the main analysis, and we summarise the other five types at the end of the results section.

**Table 1:** Traits used for comparisons of cooperative vs. private genes. The full gene lists are in supplementary tables 2-7.

| Trait | Private genes | Cooperative genes |
| --- | --- | --- |
| **1. Quorum sensing traits** | Genes which only affect the fitness of the producing cell (N=25) | Genes for public goods that potentially provide benefits to the local group of cells (N=153) |
| **2. Iron scavenging** | Genes for uptake and use of iron via the *B. subtilis* siderophore bacillibactin (N=5) | Genes for biosynthesis and secretion of bacillibactin (N=5) |
| **3. Antimicrobial resistance** | Genes for ABC transporters involved in multi-drug resistance. These genes transport the intact antibiotic outside the cell (N=11) | Genes for enzymes that deactivate beta-lactam and aminoglycoside antibiotics outside the cell (N=3) |
| **4. Proteases** | Genes for intracellular proteases, which are likely to be involved in processing and regulation of proteins within the cell (N=9) | Genes for extracellular proteases, which are likely to be involved in collective feeding and motility (N=8) |
| **5. Toxins** | Genes for contact-dependant LXG toxins, which are delivered by a Type VII secretion system (N=6) | Genes for the secreted toxin bacilysin, which is activate against a broad range of bacterial competitors (N=8) |
| **6. Antimicrobial activity** | Genes for the secondary metabolite bacillaene, which inhibits predation by *M. xanthus* (N=13) | Genes for the secondary metabolite plipistatin, which is active against fungal competitors (N=4) |

### Quorum Sensing

We started by examining genes induced by the ComQXPA quorum sensing (QS) signalling system (33, 34). This system regulates gene expression in response to the density of a diffusible signal molecule. At high cell densities, the density of the signal molecule increases, causing ComA to activate, and the upregulation of a number of traits (38–41). We categorized genes as cooperative or private based on a search of the literature in *B. subtilis* (Methods). For example, the fifteen genes coding for the exopolysaccharide EPS are classed as cooperative. EPS is the main biofilm matrix component (42), and is required for biofilm formation (43). EPS is costly to produce, it provides benefits to non-producers, and non-producers can exploit producers (44). This is a classic public good. Similarly, TasA, a protein fibre that is needed for biofilm structural integrity (45, 46), has also been shown in lab experiments to be a public good (32). Mutants lacking either EPS or TasA can also complement each other, private further evidence that the benefits of these genes are shared (47). Private genes include those coding for traits such as asparagine synthase (AsnB), which controls peptidoglycan hydrolysis for cell growth and cell-wall synthesis (48).. We found that quorum sensing controls a mixture of private and cooperative traits in *B. bacillus,* categorizing N=25 of our quorum sensing-controlled genes as cooperative, and N=153 as private (Supplementary Table 2).

We started by focusing on quorum sensing because it offers a number of advantages for our purpose. First, the large size and nature of this network means that there are sufficient private and cooperative genes for a targeted analysis (N=153 & N=25 respectively). Second, shared control by the same signalling system means that private and cooperative genes controlled by the quorum sensing system are likely to have similar levels of expression (49, 50).. Third, there is data on gene expression allowing us to directly compare expression rates (36). Fourth, the coregulation of genes acts as a control for mutations in noncoding regulatory and promoter regions that could affect the production of cooperative public goods.

### QS: Controlling for conditional expression

Differential gene expression can also influence the strength of selection and so needs to be controlled for. Theory tells us that the fraction of generations in which a trait is expressed can determine the extent to which selection is relaxed (21). To control for this, we need to compare genes that are switched on or off in the same conditions. Shared control by the same signalling system means that private and cooperative genes controlled by the quorum sensing system are likely to have similar levels of expression. We were, however, also able to test this assumption directly by examining two data sets on gene expression (36, 37). Gene expression depends strongly upon environmental conditions and so there is no single rate of gene expression for each gene.

Futo *et al*. measured gene expression at 11 different points in the formation of a biofilm, and normalised their results to the median to convert expression levels to the same scale (36). This gives us an excellent dataset to test our control for conditional expression, as we can use simple correlations between pairs of genes to see if they are up- and down-regulated at the same time. For our 178 private and cooperative genes, there are 15753 unique pairs of genes. The average correlation in gene expression for a pair of these genes is 0.302 (Spearman’s correlation). We then used a bootstrap approach to see if this correlation was greater or lower than for randomly chosen genes. We took a random set of 178 genes and calculate mean pairwise correlation in the same way, and repeated 10000 times. We found that the mean pairwise Spearman’s correlation was 0.251 for the random (bootstrap) samples, and that the correlation in expression of our quorum sensing-controlled genes was higher than 99.7% of our bootstrap samples (Figure 2). Consequently, our candidate set of genes have expression rates that are correlated significantly higher than expected by chance, supporting our choice for their use in an analysis of signature of selection (p<0.004).

**Figure 2:**
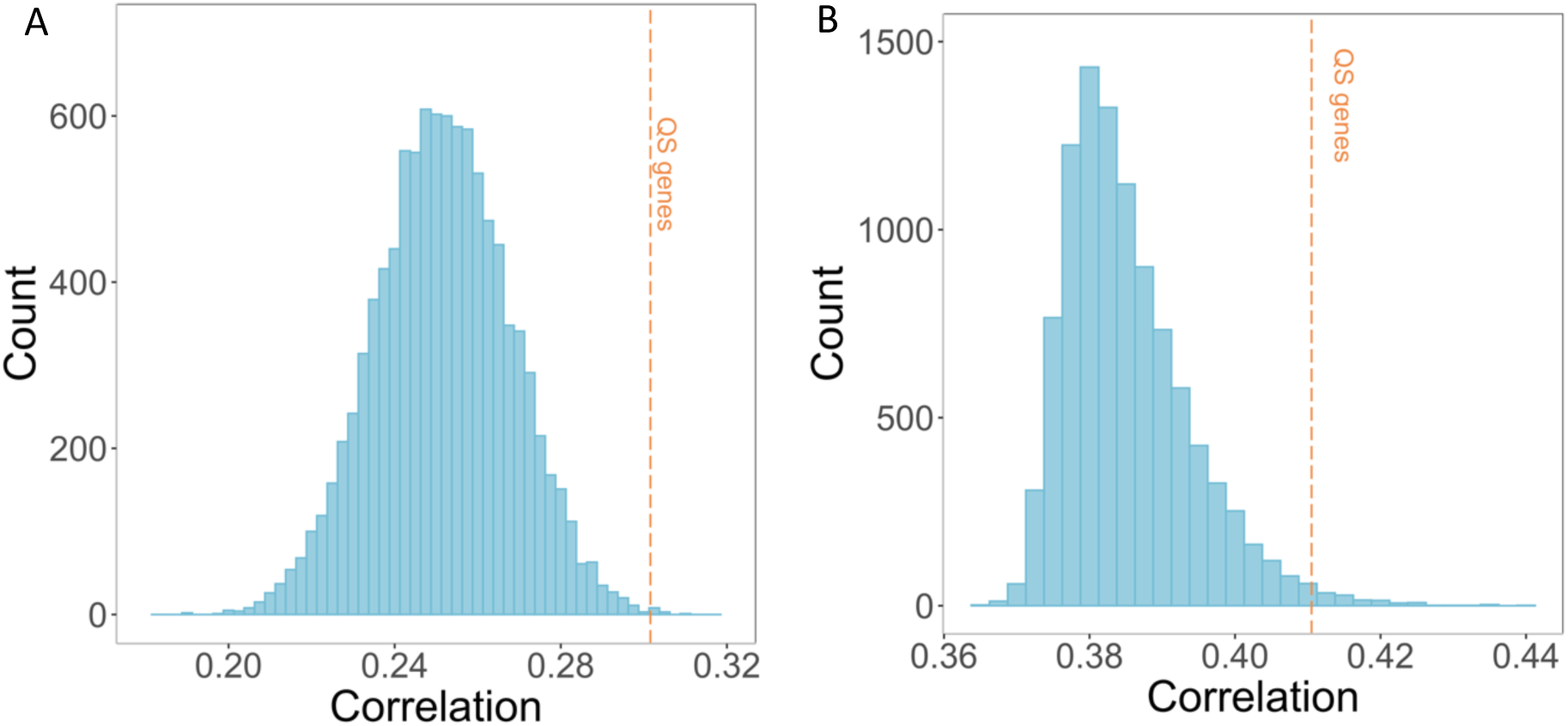
Average correlation in gene expression between genes during biofilm growth. (**A**) Correlation across 11 timepoints of biofilm formation for randomly sampled gene sets of the same size as our quorum sensing-controlled genes (N=178). Data from Futo et al. (2021). (**B**) Correlation across 8 timepoints of biofilm formation for randomly sampled gene sets of the same size as our quorum sensing-controlled genes (N=160). Data from Pisithkul et al. (2019). In both panels, the orange line shows the average correlation for the quorum sensing-controlled genes, which is >99.7% of our random samples for panel A, and >98.5% of random samples for panel B.

Pisithkul provided a different dataset measuring gene-expression in biofilms (37). Whereas Futo measured expression over two months in a biofilm with a solid-air interface, Pisithkul focussed on the initial stages of biofilm growth, measuring expression over 24 hours in a biofilm with a liquid--air interface. We again found that quorum sensing-controlled genes have expression rates that are correlated significantly higher than expected by chance (N=160 genes, p<0.02; Supplement S11).

### QS: Polymorphism

We found that genes for putatively cooperative traits had significantly greater polymorphism than genes for private traits (Figure 3; ANOVA *F*_2,61_ = 11.82, *p* < 0.0001; Games Howell Test p<0.001; N=25 cooperative genes, N=153 private genes). Cooperative genes also had significantly greater polymorphism when we only examined only non-synonymous sites (Kruskal-Wallis *X*^2^(2) = 10.7, *p* < 0.01; Dunn Test p=0.0240. Figure 4a), or only synonymous sites (ANOVA *F*_2,61_ = 7.30, *p* < 0.01; Games Howell Test p=0.007) (Figure 4b).

**Figure 3:**
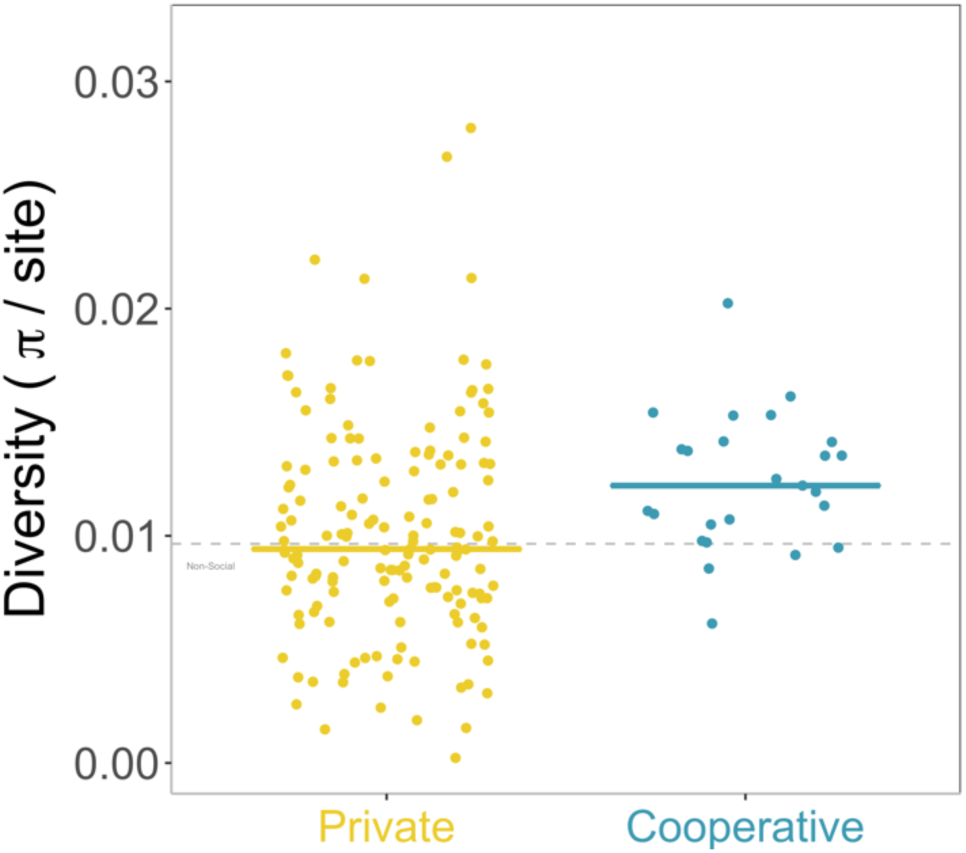
Nucleotide diversity per site for private (yellow) and cooperative (blue) genes controlled by quorum sesnsing. Each point is a gene, and the horizontal line shows the median for each group. The grey line shows the median for background private genes across the genome.

**Figure 4:**
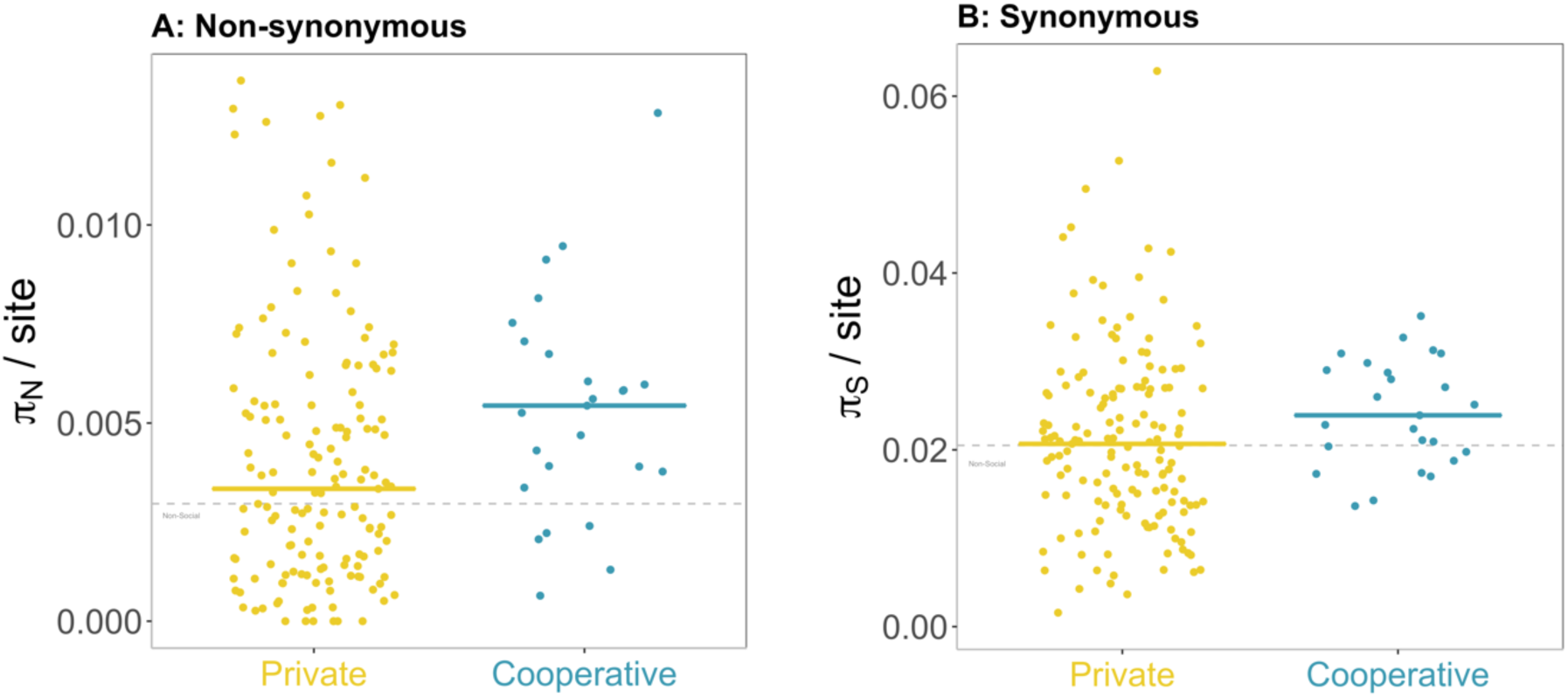
Nucleotide diversity at non-synonymous (A) and synonymous (B) sites for private (yellow) and cooperative (blue) genes controlled by quorum sensing. Each point is a gene, and the horizontal line shows the median for each group. The grey line shows the median for background private genes across the genome.

The trend for greater synonymous polymorphism is possibly surprising as such sites should be under much weaker selection, and we wouldn’t necessarily expect to see an effect of kin selection. However we also found this pattern in *P. aeruginosa* (28). There is some evidence that differences in use of preferred codons between cooperative and private genes could explain this pattern (Supplement S1). Synonymous mutations can also have substantial fitness effects on social traits (51–54).

The ratio between non-synonymous and polymorphism does not differ between cooperative and private genes (Kruskal-Wallis *X*^2^(2) = 6.97, *p* = 0.0306; Dunn Test p=0.183. Supplementary Figure 1), which reflects the fact that cooperative genes have elevated diversity at both types of site, possibly due to the large fitness effects of many traits (both cooperative and private) that are quorum sensing-controlled (55, 56) (Supplement S2).

As an additional control, we were able to repeat all of these analyses comparing cooperative quorum sensing genes against a different group of private genes not controlled by quorum sensing (N=1832). This different set of N=1832 private genes, which we call ‘background genes’ are those which aren’t controlled by quorum sensing and whose products are found in the cytoplasm, where they are least likely to have a cooperative function. In all cases we found the same pattern, with cooperative quorum-sensing genes showing elevated polymorphism compared to background genes (Supplement S2).

### QS: Divergence

We found that cooperative genes had significantly greater divergence compared to private genes at both at non-synonymous and synonymous sites (Non-synonymous: Kruskal-Wallis *X*^2^(2) = 10.4, *p* = 0.006; Dunn Test p=0.00553; Figure 5a; Synonymous: ANOVA *F*_2,61_ = 7.26, *p* < 0.01; Games Howell Test p=0.011; Figure 5b). We examined synonymous and non-synonymous sites separately because we measure divergence through rates of protein evolution. A signature of selection can be found at both synonymous and non-synonymous sites, implying that synonymous variation is not neutral (26, 28). Because divergence is elevated at both types of site (similar to polymorphism), there is no difference in the ratio of non-synonymous to synonymous divergence (Kruskal-Wallis *X*^2^(2) = 13.32, *p* = 0.00128; Dunn Test p=0.189. Supplementary Figure 2), although both sets of quorum sensing genes have a higher ratio than the background genes (Supplement S2). This could reflect stronger selection on quorum sensing traits than background traits on average.

**Figure 5:**
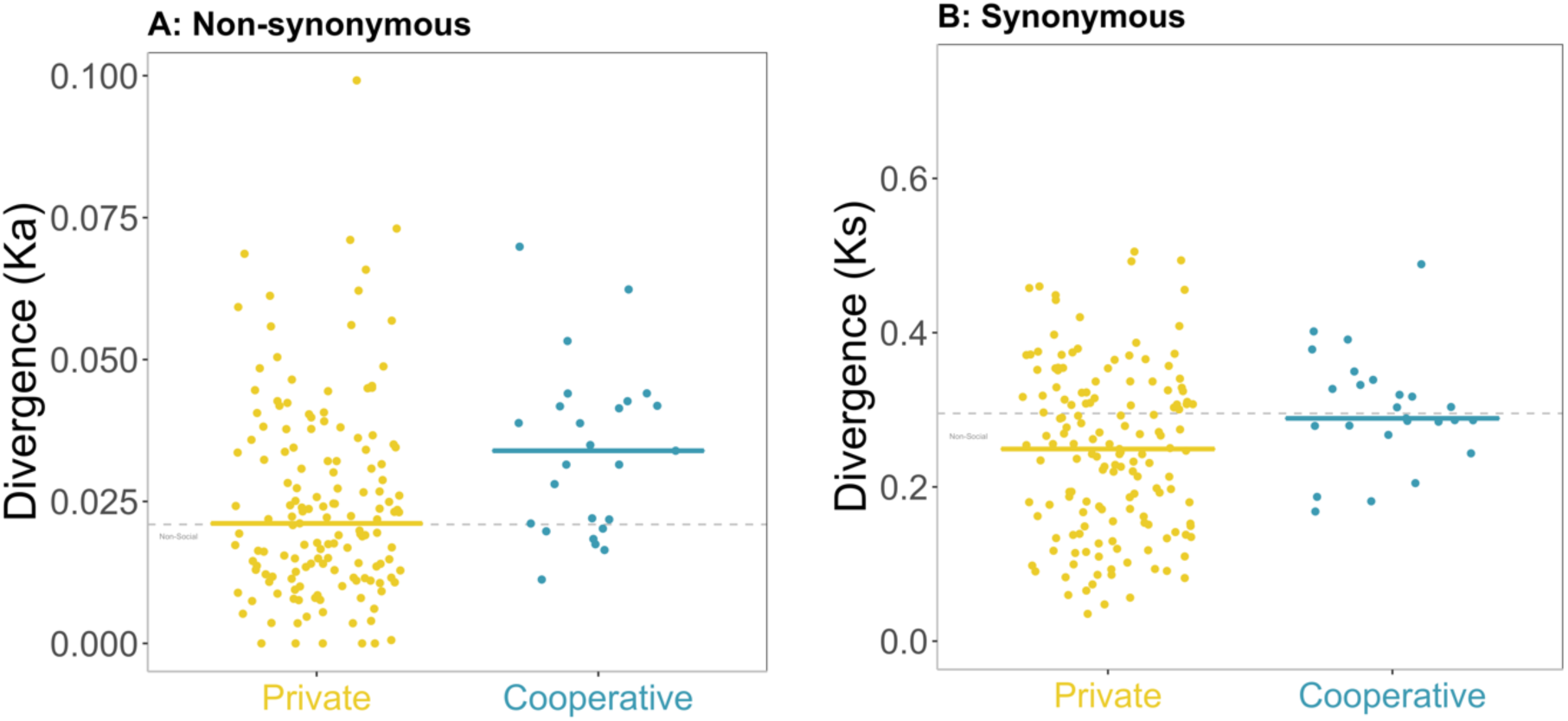
Divergence at non-synonymous (A) and synonymous (B) sites for private (yellow) and cooperative (blue) genes controlled by quorum sensing. Divergence is measured by rates of protein evolution, e.g. number of synonymous substitutions per synonymous site for panel B. Each point is a gene, and the horizontal line shows the median for each group. The grey line shows the median for background private genes across the genome.

### Alternative hypotheses

We were also able to eliminate alternative explanations for the patterns that we observed with both polymorphism and divergence (Figures 3-5). Greater polymorphism would also have been expected if cooperative genes were more likely to be experiencing balancing or frequency dependent selection, while greater divergence would arise from positive/directional selection leading to fixation of adaptive mutations (19, 20). Alternatively, the pattern we observed could have been caused by other differences between putatively cooperative and private genes. We assessed these alternative hypotheses by testing other predictions that they make.

First, we found no evidence that cooperative genes were more likely to be under balancing selection, which would lead to a deficit of rare alleles in the population (Tajima’s D, Fu & Li’s F, Fu & Li’s D, Supplement S3). Second, we found no evidence that cooperative genes are more likely to be under positive selection, which would lead to an excess of non-synonymous divergence compared to non-synonymous polymorphism, as adaptive mutations would quickly spread and only be detected as divergence (Mcdonald-Kreitman test, Neutrality Index, Direction of Selection statistic, Supplement S4). Third, we found no evidence that the patterns observed are due to any differences in gene length or likelihood of horizontal gene transfer between cooperative and private genes, or a lack of statistical power (Supplement S5). Fourth, we found no evidence that our results could be explained by noise due to variation in the recombination rate. The set of strains that we analysed vary in genetic competence, the ability to take up DNA from the environment (35), which is a form of recombination in bacteria. This variation in competency could create noise in our population genetic measures that focus on SNPs, due to the variation in recombination. Considering the 31 strains we analysed, 18 are genetically competent (35). We conducted an analysis of polymorphism and divergence using only the competent strains, and found the same patterns as when we use all strains (Supplement S7).

Finally, we found no evidence that the patterns we observed could be caused by division of labour (57–60), which is a cooperative hypothesis not mutually exclusive with kin selection. In *B. subtilis* biofilms, some cells will produce the polysaccharide EPS, and some will produce TasA amyloid fibres (61). Mutants lacking cannot grow alone but can grow together (32). The extra level of conditional expression in these public goods (over that caused by being quorum sensing-controlled), could leave a signature of relaxed selection that isn’t caused by kin selection. To investigate this possibility, we took advantage of the fact that this heterogeneity is ultimately caused by Spo0A. Spo0A is a bistable switch that is active in only a subset of cells (61, 62) and activates SinR anti-repressors which control the operons for both EPS and TasA (32, 63, 64). If the extra conditionality of division of labour was causing an effect, we would expect that genes under the control of Spo0A (N=20) would have greater polymorphism than other quorum sensing-controlled genes (N=157), but this is not the case (Supplement S6). We note that the cooperative genes EPS and TasA that are known to have division of labour both stand out as highly polymorphic within the genes controlled by Spo0A, providing support to our hypothesis that sociality causes the effect. We note that the cooperative genes EPS and TasA that are known to have division of labour both stand out as highly polymorphic within the genes controlled by Spo0A, providing support to our hypothesis that sociality causes the effect.

### Other Cooperative Traits

Our above analyses have provided strong evidence for relaxed selection due to kin selection for cooperation, considering genes controlled by quorum sensing. We then tested if the same pattern was found in five other types (groups) of traits, where we could compare genes for putatively private and cooperative traits, that are likely to be expressed at similar rates (Table 1; Figure 6). The extent to which the distinction between private and cooperative traits can be made varies across these other groups of traits. Consequently, we might not expect to see a signature of kin selection in every case, and so our aim is to see if there is a relatively consistent pattern.

**Figure 6:**
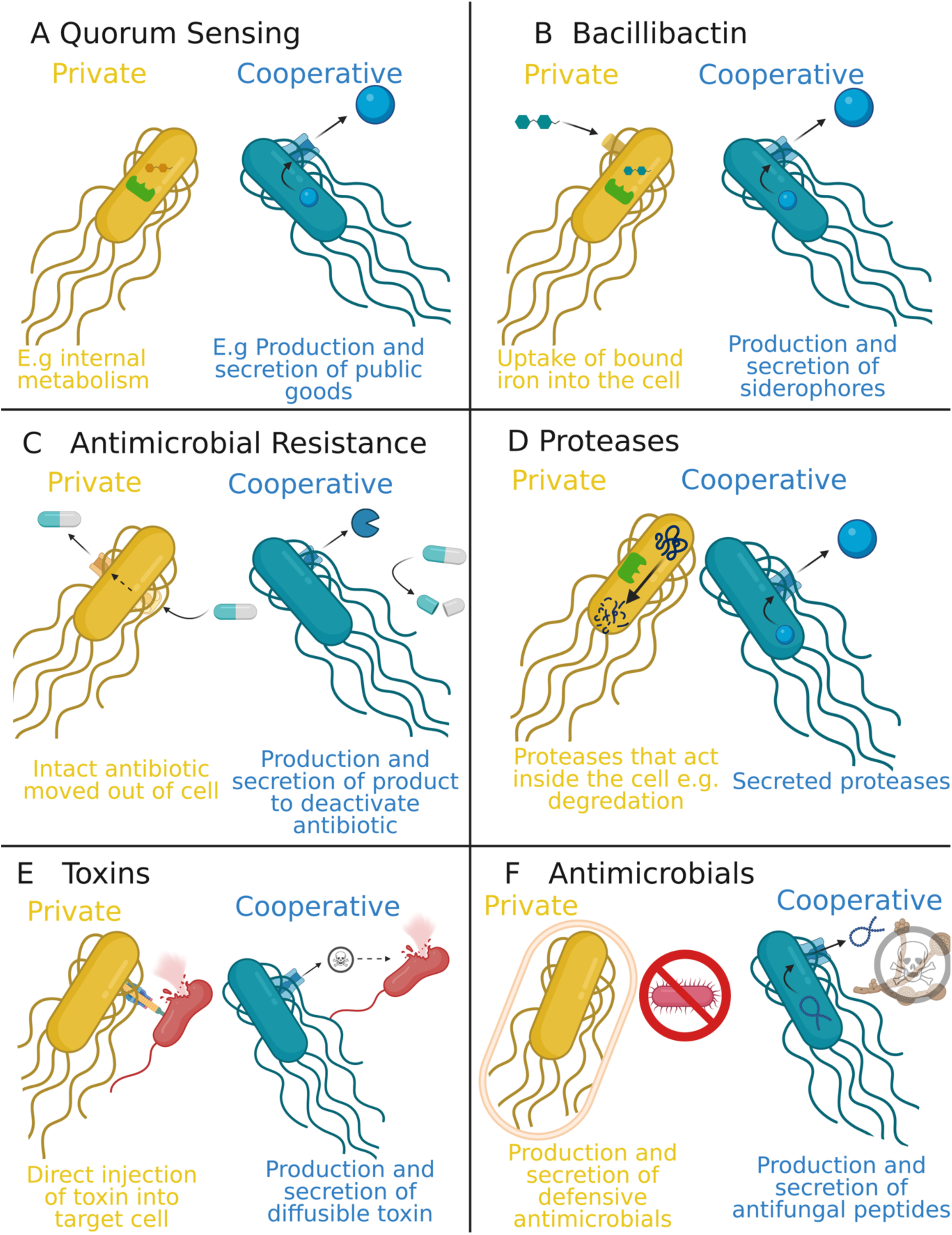
Diagram of how traits are categorised as either private (yellow) or cooperative (blue). Panel (A) shows categorisation of quorum sensing controlled traits for the main analysis, with private traits giving fitness benefits only to those expressing the gene, and cooperative traits giving fitness benefits that can potentially be shared with other cells. Panels B-F show other cooperative traits; (B) Bacillibactin (iron scavenging). (C) Antimicrobial resistance. (D) Proteases. (E) Toxins. (F) Antimicrobials. Note that for some of these comparisons it is the relative level of sociality that is different. Figure created with BioRender.com

First, *B. subtilis* produces and secretes a siderophore named bacillibactin, which binds to iron in the environment (65, 66) (Figure 6b). The bound complex can be taken into the cell, but is also available to non-producers, and is therefore a public good. We separated the genes involved in the bacillibactin pathway into cooperative and private components, with: genes involved in biosynthesis and export classed as cooperative, and those involved in uptake and release of bound iron classed as private. Second, *B. subtilis* exhibits resistance to antimicrobials by either pumping intact antibiotics outside the cell (private), or by producing enzymes such as beta-lactamases that detoxify the environment for the entire community (67–69) (cooperative; Figure 6c).

Third, *B. subtilis* produces proteases to break down proteins, with different proteases acting either inside the cell (private), or secreted to act outside the cell (70–72) (cooperative; Figure 6d). Fourth, *B. subtilis* produces toxins which can either be contact-dependant (relatively private), or diffusible throughout the community (cooperative public goods; Figure 4d). However, this comparison is relative, and possibly weak, as killing cells with contact-dependant toxins can also provide a cooperative benefit to other local cells, that experience reduced competition. Fifth, *B. subtilis* has a number of antimicrobial traits which are more defensive against predators without affecting the predators’ growth (private) and those which are more offensive against competitors (cooperative) (73, 74). This is also a weak comparison, as bacillaene (the defensive molecule) is secreted from cells, and so likely also has some cooperative component. However, the defensive traits provide a relatively more private benefit in providing personal protection, whereas the removal of competitors by the offensive traits provides a relatively more cooperative benefit. For all comparisons, we find that the set of genes have significantly correlated expression, using the same methodology and data as for the comparison with quorum sensing genes (Supplement S9) (36).

The number of cooperative genes was too small to analyse each case separately – for example, the iron-scavenging comparison involves only 10 genes (5 private and 5 cooperative). Consequently, we examined the data in three ways. First, we grouped all cooperative genes together into one set (N=52) and compared them to the grouped private genes across all comparisons (N=194). Second, we grouped all of the cooperative genes from just the five new comparisons (i.e. not quorum sensing) (N=27), and compared them to the private genes from these five comparisons (N=41). Third, we consider each of the six categories of cooperative vs. private genes (Table 1) as its own data point (N=6).

### Other Cooperative Traits: Polymorphism and divergence

We found that polymorphism was consistently higher in cooperative genes than in private genes (Figure 7). This pattern is consistent across the three different ways that we can analysed our data: all genes comparison (ANOVA *F*_2,127_ = 10.59, *p* < 0.0001; Games Howell Test p=<0.001); just the genes for the social traits other than quorum sensing (ANOVA *F*_2,47_ = 5.94, *p* < 0.01; Games Howell Test p<0.01); and the six categories comparison (Wilcoxon signed-rank test V=21, p=0.031).

**Figure 7:**
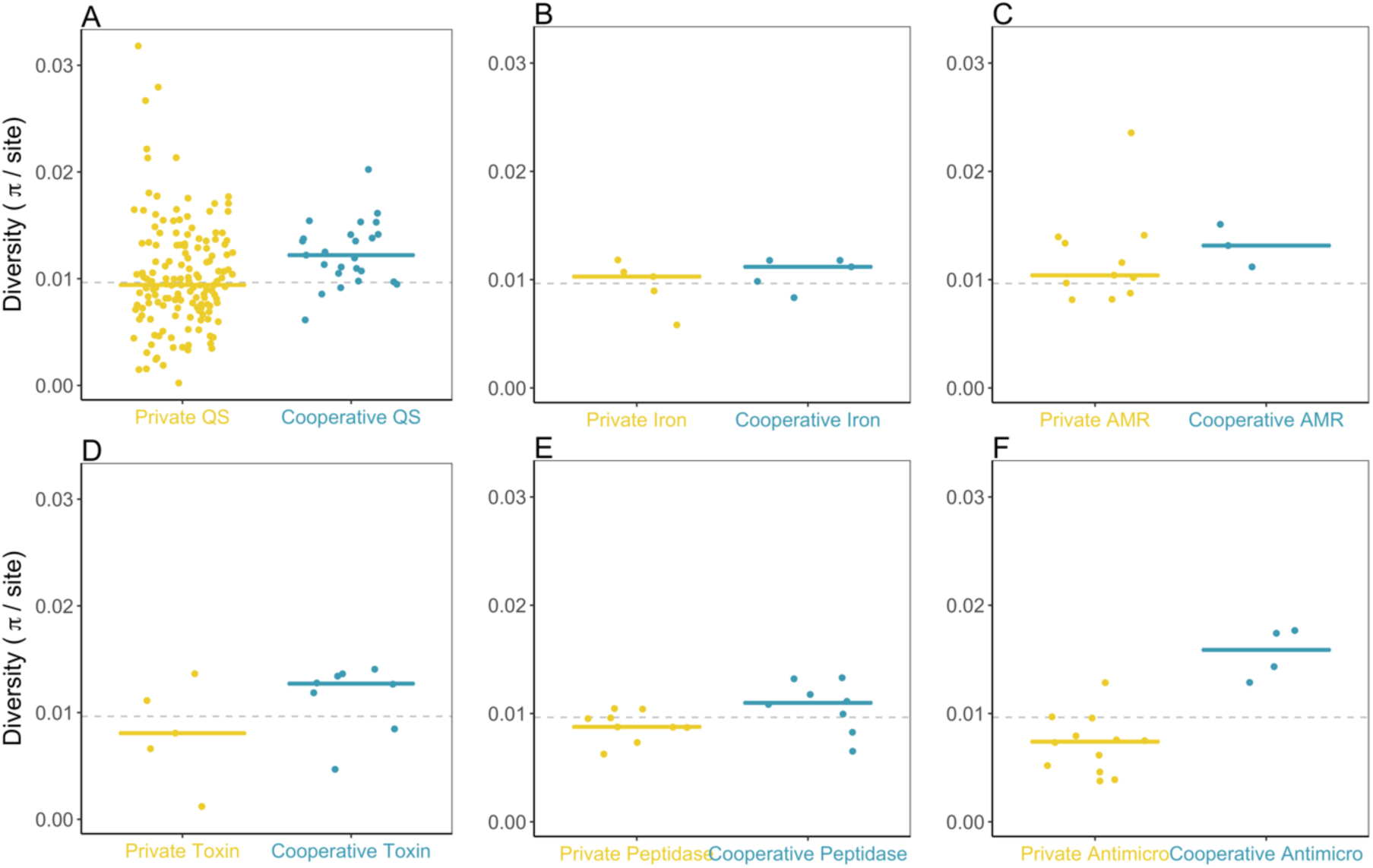
Private (yellow) vs. cooperative (blue) polymorphism in genes for six traits. Panel A shows the quorum sensing-controlled genes used in the main analysis. Panels B-F show the other cooperative traits.

Polymorphism at non-synonymous sites is also significantly higher in cooperative genes than in all private genes. This pattern is consistent across the three different ways that we can analysed our data: all genes comparison (Kruskal-Wallis *X*^2^(2) = 19.71, *p* < 10^-4^, Dunn Test *p* < 10^-4^); just the genes for the social traits other than quorum sensing (Kruskal-Wallis *X*^2^(2) = 9.90, *p* < 0.01, Dunn Test *p* < 0.01); and the six categories comparison (Wilcoxon V=21, p=0.031) (Supplementary Figure 3).

Polymorphism at synonymous sites is also significantly higher in all cooperative genes than in all private genes when analysing all genes (ANOVA *F*_2,130_( = 4.21, *p* = 0.016; Games Howell Test p=0.019). However, this trend was not significant when examining just the genes for the social traits other than quorum sensing (ANOVA *F*_2,47_ = 2.28, *p* = 0.113; Games Howell Test p=0.10), or when using categories as data points (synonymous polymorphism is marginally higher in private genes than cooperative genes for the iron-scavenging and AMR categories; Wilcoxon V=18, p=0.156). We would expect the pattern to be weaker with synonymous polymorphism, as these sites are likely to be under weaker selection.

Non-synonymous divergence is significantly greater in all cooperative genes compared to all private genes. This pattern is consistent across the three different ways that we can analysed our data: all genes comparison (Kruskal-Wallis *X*^2^(2) = 22.9, p < 10^-4^, Dunn Test p < 10^-4^); the genes for the social traits other than quorum sensing (Wilcoxon V=21, p=0.031 Supplementary Figure 5); and the six categories comparison (Kruskal-Wallis *X*^2^(2) = 14.7, *p* < 0.001, Dunn Test p < 0.001). As would be expected, the pattern is more mixed for synonymous divergence. Whilst synonymous divergence is significantly greater in all cooperative genes compared to all private genes (ANOVA *F*_2,125_ = 8.33, *p* < 0.001; Games Howell Test p<0.01), this isn’t the case when examining just the genes for the social traits other than quorum sensing (ANOVA *F*_2,45_ = 2.10, *p* = 0.13; Games Howell Test p=0.39), or the six categories comparison (Wilcoxon V=13, p=0.688) (Supplementary Figure 6). Similarly, the ratio between non-synonymous and synonymous divergence is significantly greater in all cooperative genes compared to all private genes (Kruskal-Wallis *X*^2^(2) = 25.0, < 10^-5^, Dunn Test p = 0.021), but not when we just looked at the genes for the social traits other than quorum sensing (Kruskal- Wallis *X*^2^(2) = 13.2, < 0.01, Dunn Test p = 0.11) or the six categories comparison (Wilcoxon V=18, p=0.156) (Supplementary Figure 7). This may reflect differences in selection on synonymous variation on different traits, or the weakness of some of these comparisons due to small sample sizes. We confirmed that none of our cooperative vs. private comparisons show significant differences in balancing or positive selection (Supplement S10), suggesting that what we are seeing is a signature of kin selection, and that we may just lack power in our other comparisons.

Overall, these results incorporating other traits in addition to quorum sensing controlled genes, provide support to the main result that there is a signature of kin selection for cooperation (Figures 3-5). Nonetheless, the *a priori* distinction between private and cooperative traits is weaker for some comparisons, and we could expect exceptions within the overall pattern. The main exception in our analyses was toxin comparison, where we compared contact-dependant LXG toxins (N= 6 genes) to the secreted antimicrobial bacilycin (N=8 genes). Both of these sets of genes are involved in competition in biofilms, and both are controlled by the DegS-DegU system, so likely expressed at similar rates (75, 76). Possible confounding factors in this case include the fact that although we classified the contact-dependent toxins as private, they also provide cooperative benefits to local cells by eliminating competitors. The LXG toxins also stand out because three of them are on phage elements (75), and it may be that the strength or type of selection is different on these genes, masking any effect of sociality.

### All traits: Deleterious mutations

As an additional robustness test for our conclusions, we analysed deleterious mutations. If kin selection is favouring cooperation, we should also observe more deleterious mutations in genes controlling cooperative traits compared to private traits. This is because relaxed selection slows the rate at which deleterious mutations are purged from the population (19, 20, 22). We tested this prediction by looking for loss-of-function mutations, that generate stop codons or frameshift mutations, and hence act as deleterious mutations, in our SNP data. We repeated this analysis with two different data sets.

First, we used all cooperative genes from our six comparisons in Table 1. We measured how many cooperative and private genes have deleterious mutations, and compared this to an expectation based on their relative frequency across the genome. Cooperative genes were significantly more likely than private genes to have deleterious mutations (X^2^ (1) = 12.3, *p* < 0.001. This pattern also holds if we count total number of deleterious mutations, rather than just presence or absence (X^2^ (1) = 11.0, *p* < 0.001.

To test the robustness of this result, we repeated the analysis using the localization prediction tool PSORTb to categorize genes for extracellular proteins as ‘cooperative’, and genes for proteins that aren’t secreted as ‘private (77). This method has been previously used in several studies to estimate whether genes are for cooperative (social) or private traits (78, 79). By using PSORTb we are able to analyse all genes, which increases our sample size and statistical power. We removed the 17% of all genes with unknown localization, leaving us with a set of genes of known sociality. We found deleterious mutations in 293 genes, of which 17 are cooperative (5.8%). This is significantly more than expected given that cooperative genes only make up 2.0% of genes (binomial test P<0.001), matching our prediction that deleterious mutations should be biased towards cooperative genes. If we count total deleterious mutations (rather than number of genes with at least one) we see the same pattern, with cooperative genes making up 5.5% of mutations (20 of 361).

### Relatedness estimation

The genetic relatedness between interacting cells (*r*) is a key parameter for social evolution. Relatedness can be very hard to estimate for natural populations of bacteria and other microbes, except for extreme cases where interactions take place in some physical structure such as a fruiting body or a filament (80, 81). Population genetic data allows relatedness to be estimated indirectly, because the degree to which selection is relaxed, and greater polymorphism will be observed, depends upon the relatedness between interacting cells. Consequently, if we assume that the patterns of polymorphism and divergence that we see are due to relatedness being <1, we can work backwards from the polymorphism data to obtain an indirect estimate of relatedness (28). Examining the polymorphism data from all genes, we estimated relatedness to be r=0.77 (95% CI 0.67-0.91) (Supplement S8). This estimate assumes that both the magnitude of selection and the distribution of selection coefficients is the same on average for cooperative and private genes. An advantage of this indirect method for estimating relatedness is that it does not require knowledge about factors that would be hard or impossible to measure in natural populations. For example, the spatial details of how cooperative interactions play out, such as how far do public goods diffuse, and who benefits, as well as how much these vary in different environments, and the frequency with which different environments are encountered (28, 82). Indeed, cooperative traits in *B. subtilis* vary in the degree to which they are shared depending on whether groups are exhibiting sliding motility or growing in biofilms (16). In contrast, the indirect measure provided by population genetics represents an average for the different cooperative traits, over the different environments encountered, over evolutionary time.

## Conclusions

We have found strong evidence of kin selection for cooperation in a natural population of *B. subtilis.* Our analyses controlled for possible confounding factors, such as expression rate, and eliminated alternative explanations for polymorphism and divergence, providing evidence that complements the lab experiments demonstrating sociality in this species (32, 42–46, 61, 83– 86). Taken together with a previous study, population genetic analyses have now provided evidence of kin selection for cooperation in both a gram-positive (*B. subtilis*) and a gram-negative (*P. aeruginosa*) bacteria (28). These results suggest convergent, and potentially widespread, kin selection for cooperation, based on very different underlying mechanisms, across bacteria.

A possible complication with studying cooperation in bacteria is that the extent to which traits are cooperative, and their importance for fitness, can depend greatly upon environmental conditions (8, 87–90). One advantage of the molecular population genetic approach that we have used is that it averages across different environments over evolutionary time. Consequently, rather than examining a specific environment, it provides an ‘average’ answer. In the case of *B. subtilis,* for the traits that we have examined, we have found evidence of kin selection for cooperation, with an estimated average relatedness of *r*=0.77.

## Acknowledgements

We thank: Ming Liu and Ákos Kovács for useful discussion and comments on the manuscript. This work was supported by the European Research Council (834164: LJB, AED & SAW; SESE: MG).

## Material & Methods

### Strains

We use the whole-genome sequences of 31 strains of *Bacillus subtilis* from (35). The strains are environmental isolates, collected from a citizen science project in Dundee where people brought soil samples from their garden. *B. subtilis* is most commonly found in soil, but is also found living as a commensal in animal intestines (91), and in marine environments (92).. Whilst durable spores than can disperse through the air can allow long-distance migration (93), and strains don’t phylogenetically cluster based on environment (94), several factors are in favour of using these strains as our natural population.. Firstly, these samples were all collected at the same time for the same project (35). Further, we know that rates of migration between populations scales with geographic distance, most of the sequence diversity within the species in contained in local population (93), and there is evidence for fine-scale genetic structure in micro-scale populations (95).

We downloaded raw sequence data for each strain from the European Nucleotide Archive (accession number PRJEB43128). The full list of strains can be found in Supplementary Table S1.

**Supplementary Table 1:**
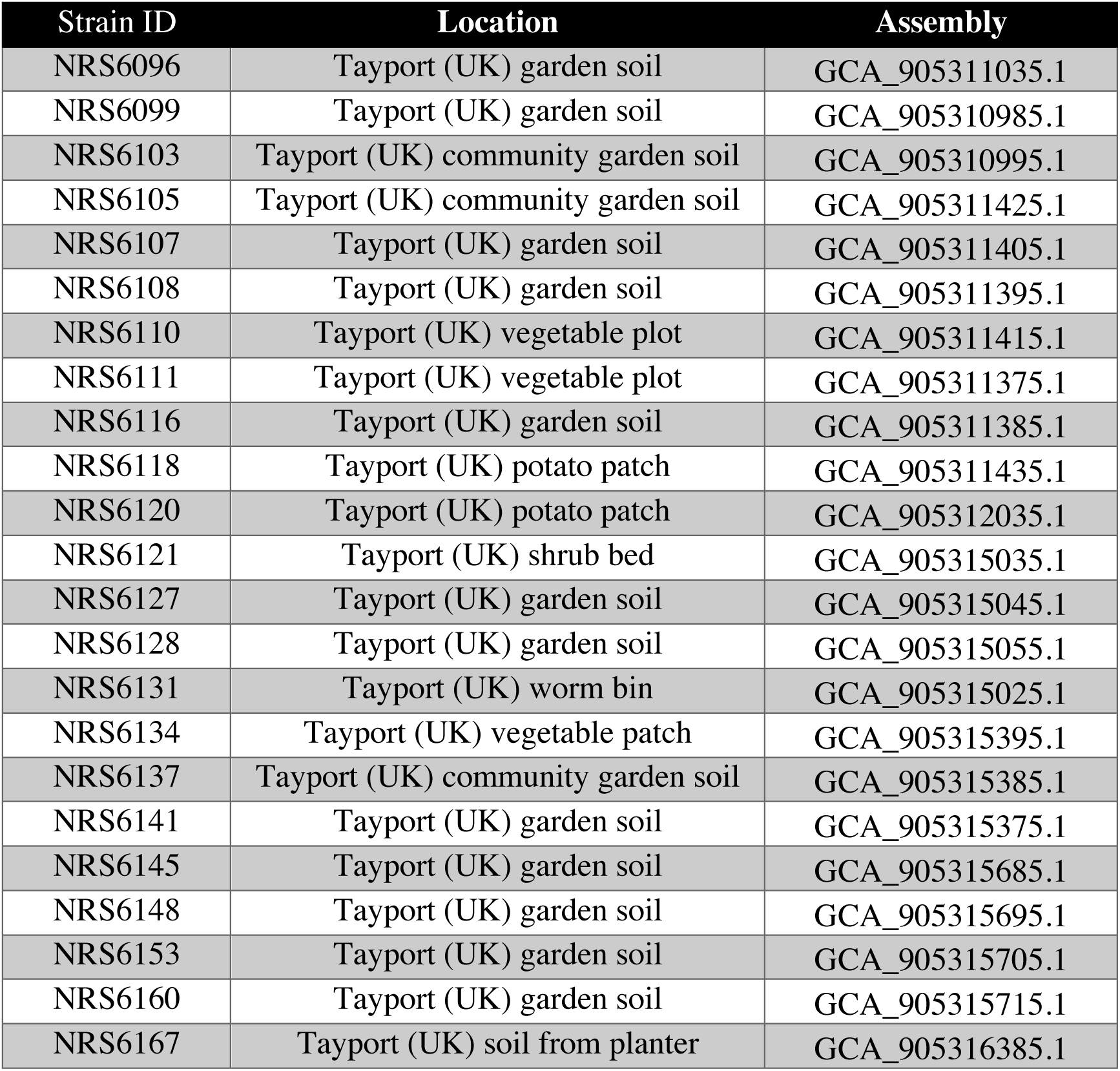
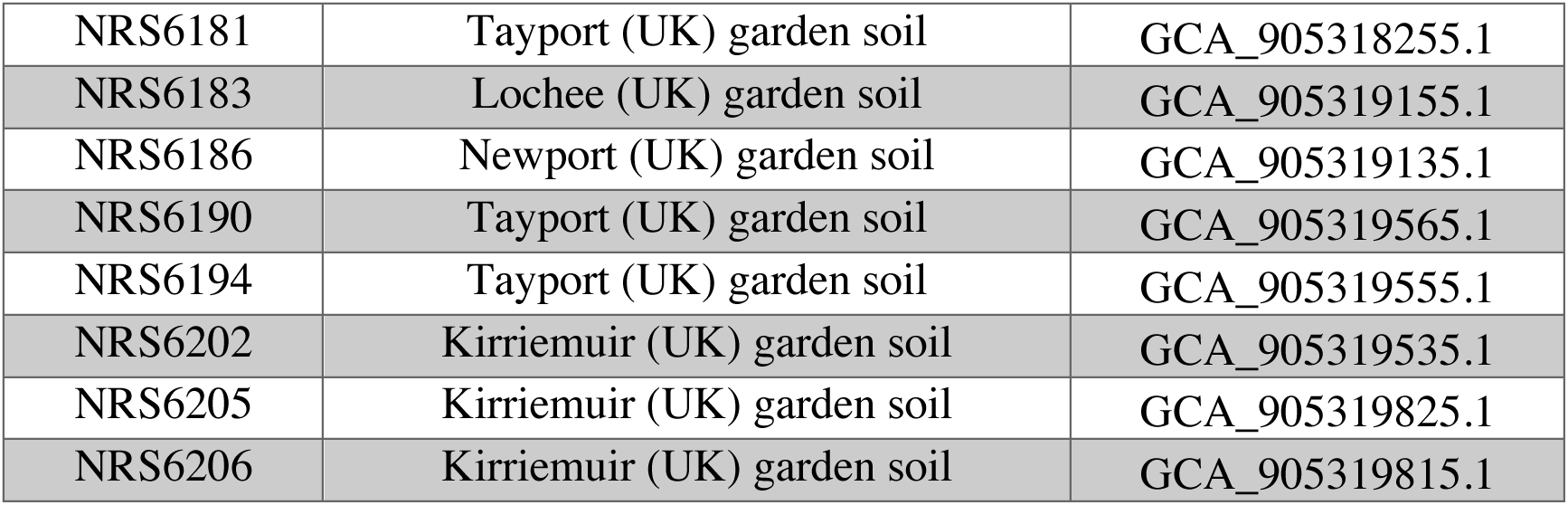
List of strains used

### Genes regulated by quorum sensing

For our set of quorum sensing controlled genes, we combine three published datasets: (1) 88 genes controlled by ComXAP (40); (2) 114 genes controlled by degU (96); (3) 40 genes controlled by Spo0A (63).. We didn’t use a fourth possible dataset, of 166 genes affected by the competence transcription factor ComK, which are indirectly regulated by ComA (97).. This is partly because our undomesticated reference strain (NCIB 3610) carries a plasmid-encoded protein which interferes with the competence machinery (98) (Supplement S7).). In addition, we wanted to focus on the quorum sensing systems known to produce public goods (Figure 3 of (34)).

### Identifying social genes

We used an artisan approach to identify social genes based on laboratory studies which have demonstrated that a trait is cooperative. The gold-standard test for a cooperative gene involves a wildtype strain which produces the traits, and a mutant strain which doesn’t. If a trait is cooperative, then the producer will outperform the non-producer when each is grown clonally, but non-producers will outperform producers in groups (2). As an example, we look at the first gene on the list, bslA (formely known as yuaB), which is involved in biofilm formation, and specifically in making the biofilm hydrophobic to resist chemical attack (99). A non-producer of bslA cannot form normal biofilms on its own, but can get into mature biofilms when in co-culture with producers (100). Further work mixing producers and non-producers at a range of starting ratios demonstrated that the biofilm can maintain function as long as >50% of cells are producers (99).

The full list of cooperative genes can be found in Supplementary Table 2.

**Supplementary Table 2:** List of social genes

| Gene | Function | Reference |
| --- | --- | --- |
| ***bslA*** | biofilm-surface layer protein BslA | Arnaouteli 2020 |
| ***bslB*** | biofilm-surface layer protein BslB | Morris 2017 |
| ***epsA*** | hypothetical protein | Arnaouteli 2020 |
| ***epsB*** | protein tyrosine kinase EpsB | Arnaouteli 2020 |
| ***epsC*** | polysaccharide biosynthesis protein | Arnaouteli 2020 |
| ***epsD*** | glycosyltransferase family 4 protein | Arnaouteli 2020 |
| ***epsE*** | glycosyltransferase EpsE | Arnaouteli 2020 |
| ***epsF*** | glycosyltransferase family 1 protein | Arnaouteli 2020 |
| ***epsG*** | biofilm exopolysaccharide biosynthesis protein EpsG | Arnaouteli 2020 |
| **epsH** | glycosyltransferase | Arnaouteli 2020 |
| **epsI** | polysaccharide pyruvyl transferase family protein | Arnaouteli 2020 |
| **epsJ** | lipoprotein | Arnaouteli 2020 |
| **epsK** | cyclic-di-GMP receptor EpsK | Arnaouteli 2020 |
| **epsL** | sugar transferase | Arnaouteli 2020 |
| **epsM** | acetyltransferase | Arnaouteli 2020 |
| **epsN** | aminotransferase class I/II-fold pyridoxal phosphate-dependent enzyme | Arnaouteli 2020 |
| **epsO** | polysaccharide pyruvyl transferase family protein | Arnaouteli 2020 |
| **sipW** | signal peptidase I | Arnaouteli 2020 |
| **srfAA** | surfactin non-ribosomal peptide synthetase SrfAA | Kalamara 2018 |
| **srfAB** | surfactin non-ribosomal peptide synthetase SrfAB | Kalamara 2018 |
| **srfAC** | surfactin non-ribosomal peptide synthetase SrfAC | Kalamara 2018 |
| **srfAD** | surfactin biosynthesis thioesterase SrfAD | Kalamara 2018 |
| **srfT** | MFS transporter | Kalamara 2018 |
| **tapA** | amyloid fiber anchoring/assembly protein TapA | Arnaouteli 2020 |
| **tasA** | biofilm matrix protein TasA | Arnaouteli 2020 |

For the robustness check of whether deleterious mutations are over- or under-represented in cooperative genes, we used the protein localisation tool PSORTb 3.0 (77). We categorize cooperative genes as those which PSORTb predicts to be extracellular. We also follow previous studies in removing genes for which PSORTb cannot make a definitive prediction (78)

### Controlling for conditional expression

Conditional expression can lead to the same signatures of relaxed selection as kin selection for cooperation. We directly examined expression rates for the set of genes regulated by quorum sensing, using data from (36), who measured gene expression of >4000 genes at 11 timepoints during biofilm formation.

For any pair of genes, we can calculate the correlation in gene expression across the 11 timepoints of the biofilm. For the 178 quorum sensing genes in our dataset, there are 15753 unique pairs of genes. The mean pairwise Spearman’s correlation in gene expression is 0.302. To test whether this set of genes is more or less correlated than a randomly chosen set of genes, we use a bootstrap approach. We take a random set of 178 genes and calculate mean pairwise correlation in the same way as before. Then we repeat 10000 times. We find that the correlation in expression of our quorum sensing-controlled genes is higher than in 99.7% of our bootstrap samples (Supplementary Figure 3), demonstrating that our candidate set of genes is appropriate for our analysis of signature of selection.

### Other cooperative traits

We also examined five other types (groups) of traits, where we could compare genes for putatively private and cooperative traits, that are likely to be expressed at similar rates (Table 1; Figure 4). Firstly, we used iron-scavenging via siderophores, which is a well-studied cooperative trait that is important for growth and survival of bacteria (3, 101). Specifically, we looked at the *B. subtilis* siderophore bacillibactin (65, 66). We classified the genes for biosynthesis bacillibactin as cooperative, and genes for uptake and release of bound bacillibactin as private (Supplementary Table 3).We classified the genes for biosynthesis bacillibactin as cooperative, and genes for uptake and release of bound bacillibactin as private (Supplementary Table 3).

Second, we looked at antibiotic resistance genes. There are many mechanisms of antibiotic resistance, some of which are cooperative. For example, the secretion of beta-lactamases is a cooperative trait as they detoxify the external environment, providing benefits to the local population (5, 102). We also classified aminoglycoside resistance as cooperative, as the modification of the antibiotic detoxifies the local environment (103). For the private genes, we used the eight ABC transporters that are thought to be involved in multi-drug resistance by pumping antibiotics outside the cell (69, 104) (Supplementary Table 4).

Thirdly, we looked at the range of peptidase proteases produced by *B subtilis*. The functions of these proteases are broad, covering processing, regulation, and feeding, but we can separate them in cooperative and private genes by looking at those which are secreted (i.e. are extracellular) and those which are not secreted (70). The secreted proteases are more likely to have cooperative fitness effects on other cells, through nutrition, interacting with host immune systems etc (Supplementary Table 6)..

Fourthly, we looked at toxins. For the cooperative genes, we used bacilycin, which is a secreted antimicrobial peptide that is active against a range of bacteria (105). Because bacilycin can diffuse through the environment, it likely has cooperative fitness effects on others. *B. subtilis* also has many toxins which are involved in contact-dependent inhibition, and therefore likely have private effects on fitness. For the private gene, we used six LXG toxins (75), which are delivered by a Type VII secretion system. Whilst these toxins can still have cooperative effects by removing competitors, they are by their nature less cooperative than secreted molecule such as bacilycin. Both of these sets of genes are under control of the DegS-DegU system (Supplementary Table 5).

Fifthly, we looked at antimicrobials. *B. subtilis* produce a series of antimicrobial molecules, which vary in which organisms they target, how they act, and how they are secreted. We can however distinguish between antimicrobials that have a more defensive role in traits such as predation avoidance, and those which have a more offensive role in competition with other species. Whilst both of these categories likely have some component of cooperative and private effects on fitness, the offensive ones will be relatively more cooperative. This is because defensive molecules have a stronger effect on the individuals producing them (Supplementary Table 7).

### Identifying deleterious mutations

We used the variant annotation tool SnpEff (106) to look for SNPs that generate deleterious mutations in our dataset.. Specifically, we annotate two types of mutations; (1) premature stop codons and (2) frameshift mutations. This gives us a list of genes that have at least one deleterious mutation. To test if a given set of genes are overrepresented for deleterious mutations, we use two percentages; (1) the % of genes in the whole genome that are in that set, and (2) the % of genes with deleterious mutations that are in this set. We compare these values using a binomial test with the null hypothesis that the number of deleterious mutations in the gene set is equal to that expected by the frequency of the gene set. For any given gene set, we conduct a further test where we use the total number of deleterious mutations in that set of genes, rather than just presence/absence of deleterious mutations for that gene.

### Statistics & Figures

We conducted all statistical analysis in R (107).. For the main statistical analysis comparing molecular population genetic parameters of cooperative and private genes we use one of two statistical tests, depending on the variable in question. For variables that are normally distributed, we used an ANOVA to compare the three groups of genes. Because of unequal sample sizes we used Welch ANOVA which doesn’t require equal variance. We used the Games-Howell post-hoc test, which is similar to Tukey’s HSD, but designed for Welch ANOVA where we don’t have to assume equal variance. For variables that aren’t normally distributed, we used the Kruskal-Wallis test, which compares medians. We then used the Dunn test for post-hoc comparisons of groups.

All results figures were made using the ggplot2 package in R (108) using colour palettes from the packages wesanderson (github.com/karthik/wesanderson) and BirdBrewer (https:///github.com/lauriebelch/BirdBrewer). Figure 4 illustrating the secondary comparisons was made using BioRenderFigure 4 illustrating the secondary comparisons was made using BioRender.

### Bioinformatics

Raw reads for each of the 31 strains were downloaded from the European Bioinformatics Institute’s European Nucleotide Archive (accession number PRJEB43128). We then used a SNP calling pipeline to find SNPs in each strain compared to the reference NCIB 3610 (accession NZ_CP020102.1).

#### Trimming and quality-control

We used Trimmomatic to remove adapters remove low quality reads, which we did by remove leading and trailing reads if the quality score was <3 or if average quality in a four-base sliding window was <20. We manually checked the output of this step using the reports produced by FastQC (109).

#### Mapping

We used BWA (110) to map reads from each strain to the reference strain. We used SAMtools (111) to convert the mapping files from BAM to SAM, and used Picard tools (112)to remove PCR duplicates.

#### Variant calling

We used BCFtools (113) to call variants on all strains and produce a VCF file that can be read by R for population genetic analysis.

#### Filtering and quality control

We conducted further filtering to remove indels, and filter for mapping quality, read depth, and strain bias using the default setting of SAMtools vcfutils python script. We then removed all sites which hadn’t been called in at least 80% of strains. We also used the coverageBed tool in BEDtools (114) to record the percentage of each genes length that had been mapped, in order to adjust per-site measures to the correct length. After filtering, we had a total of 256,769 SNPs among the 31 strains.

#### Outgroup

We used the phylogeny in Kalamara 2019 (the source of these strains) to identify *Bacillus subtilis subsp. spizizenii* str. W23 as an appropriate outgroup (accession NC_014479.1, raw sequencing data SRR2063059). We used the same variant calling pipeline as above to produce a second VCF which included the SNPs from the outgroup.

### Population genetic measures

We used the PopGenome package from R (115) to conduct the main molecular population genetic analysis. All parameters were scaled to the corresponding mapped gene length, and any gene with mapped length <50% of their full length or lacking polymorphism data was removed from the analysis, leaving 3817 genes for the population genetics analysis. Using PopGenome, we calculated Nucleotide polymorphism, Tajima’s D, Fu and Li’s D*, the Mcdonald-Kreitman p-value, Direction of Selection statistic, and neutrality index. We also calculated separate measures for synonymous and non-synonymous sites where appropriate.

To calculate divergence, we measured the rate of protein evolution K_a_/K_s_ by comparing the reference strain to the outgroup. We did this by creating a pseudo-genome of the outgroup by inserting the relevant SNPs into the reference sequence using the GATK suite of tools (116). This pseudo-genome could then be read by R, and we used the seqinR package (117) to calculate divergence.

## Supplementary Material

### S1: Codon usage

In the main text, we showed that the elevated polymorphism in cooperative genes relative to private genes occurs at both synonymous and non-synonymous sites. This may be due to selection on synonymous codon usage, which is common across bacteria (118). In general, some codons may be preferred due to GC or AT preference, tRNA availability, or metabolic costs (119–122). We investigated whether codon usage differences between cooperative and private genes can explain the elevated synonymous polymorphism observed in social genes.

We used the R package ‘sscu’ (strength of selected codon usage), which calculates several measures of codon usage bias (123).

For many measures, we compare social genes to a set of highly expressed genes, under the assumption that highly expressed genes are under the strongest selection to optimize codon usage. We followed Rocha 2004 (122) in choosing genes that code for ribosomal proteins as this set. For our social genes, we used the set of genes coding for extracellular proteins (as determined by PSORTb), as this is a measure that can be systematically applied to the whole genome (unlike our ‘artisan’ method, which requires manual curation of genes).

The first measure we calculated is RSCU, relative synonymous codon usage. This gives a measure of the relative use of each codon, compared to the null expectation that each codon for an amino acid is used equally (RSCU =1) (Supplementary Figure S1.1).

**Supplementary Figure S1.1:**
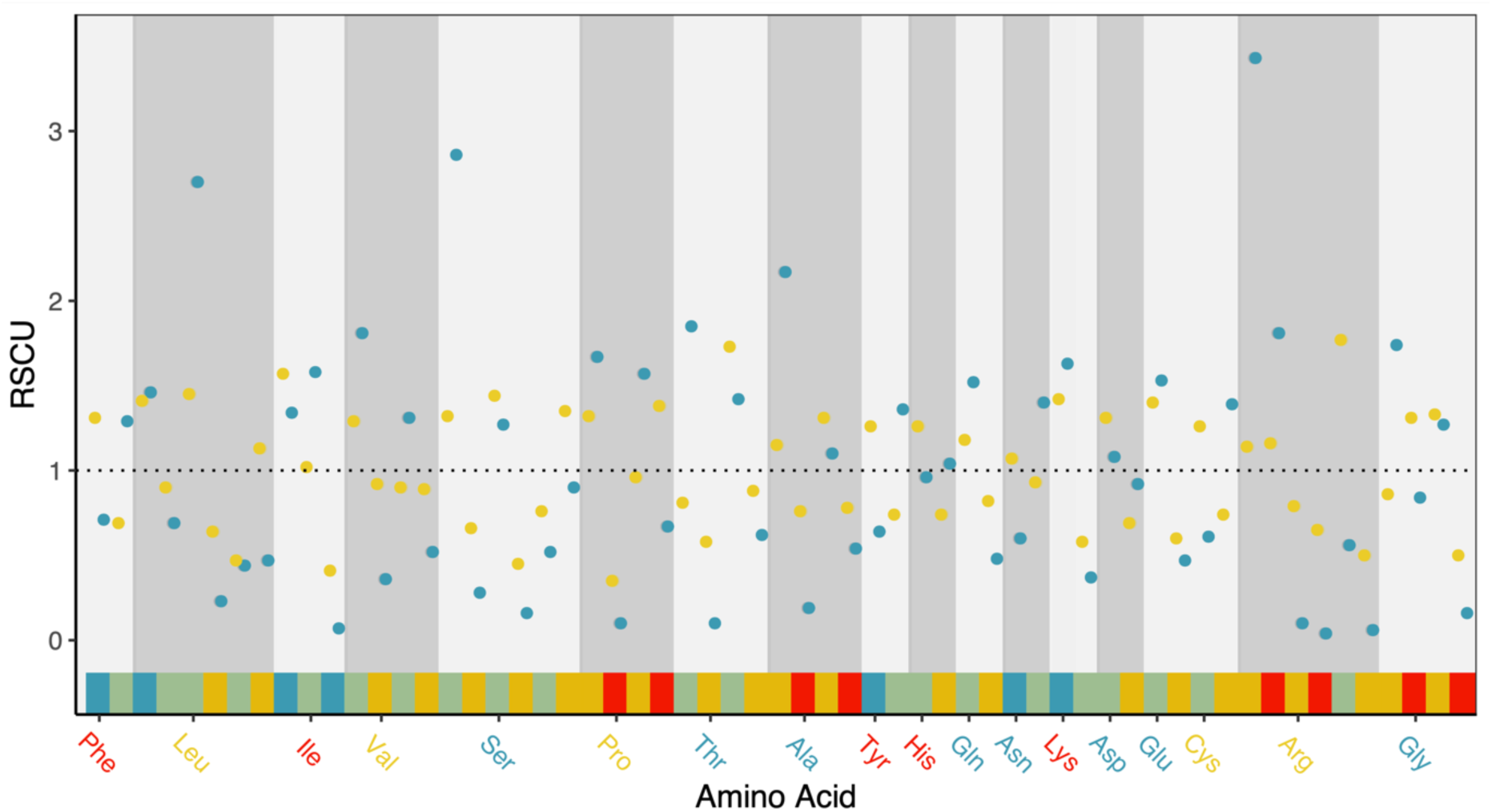
RSCU for cooperative genes (blue) alongside highly expressed (ribosomal) genes (yellow). The dotted line shows RSCU=1, which is when each codon for an amino acid is equally likely to be used. If codons are used at random, we would expect all RSCU values to be close to 1. If certain codons are strongly preferred, then each amino acid should have a codon for which RSCU>>1

To determine if codon usage is more even in social genes or highly expressed genes, we calculate Shannon entropy for each amino acid. If each codon is used evenly, then Shannon entropy will be 0. If one codon is used more frequently than random, then Shannon entropy will be negative. This is analogous to how entropy is used in ecology to calculate species richness. An amino acid mostly being coded for with the same codon is equivalent to one species dominating in a species richness measure.

On average, Shannon entropy is much less negative for cooperative genes (-0.215 compared to -0.869 for highly expressed genes), suggesting that cooperative genes have much more even usage of codons, suggesting that there may be relaxed selection for synonymous codon usage in cooperative genes (Supplementary Figure S1.2)

**Supplementary Figure S1.2:**
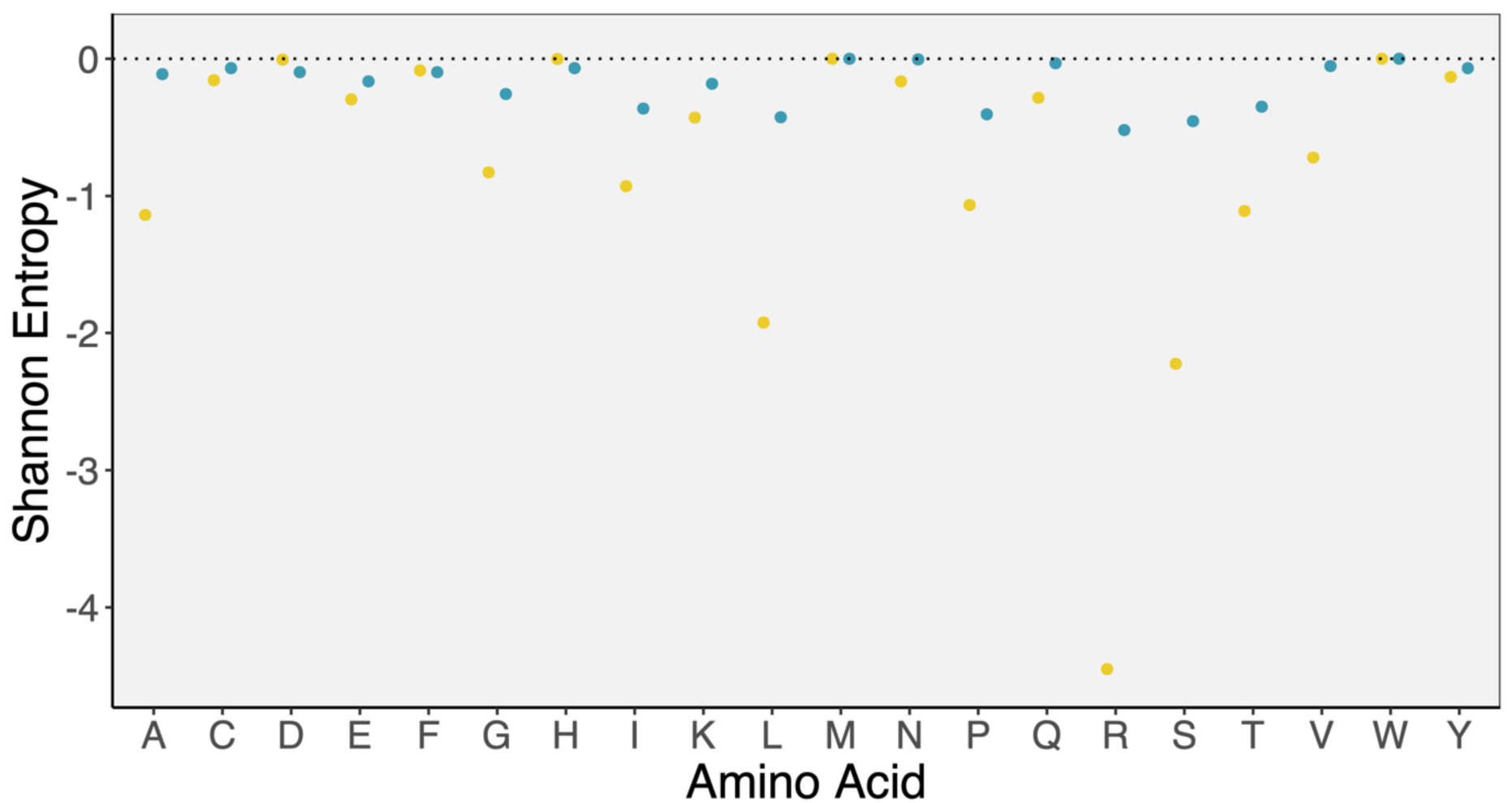
Shannon entropy for cooperative (blue) and highly-expressed (yellow) genes for each amino acid.

Cooperative genes also have higher GC3 (GC content at third positions) than our set of highly expressed genes. [0.407 in cooperative compared to 0.322 in ribosomal] which may relate to variable expression.

The effective number of codons is also substantially higher in cooperative genes than in our set of highly expressed genes (55.1 compared to 43.4), further demonstrating how cooperative genes are using more codons, and therefore less likely to use preferred codons. We also conducted chi-squared tests to determine for each codon whether it was used significantly more than expected (from random chance) in cooperative genes than highly expressed genes, and found that only 14 codons are used significantly more often than expected by chance in cooperative genes, compared to 22 codons that are used significantly more often than expected for highly-expressed genes.

Overall, there is some evidence that decreased usage of preferred codons may explain the increase in synonymous polymorphism. This is the third study to find the counter-intuitive pattern of increased synonymous polymorphism in social genes (*P. aeruginosa* (28) ; *D. discoideum* (26), so more research is needed in this area.

### S2: Cooperative vs. background genes

We compared cooperative genes to private genes in the main analysis, and here we also compare to a set of background genes. For the background genes we use those which produce proteins that localise to the cytoplasm, as these are least likely to have cooperative functions. This produces a set of 1832 genes.

Here, we present the results of the post-hoc comparison between cooperative genes and background genes for the main set of molecular population genetic measures

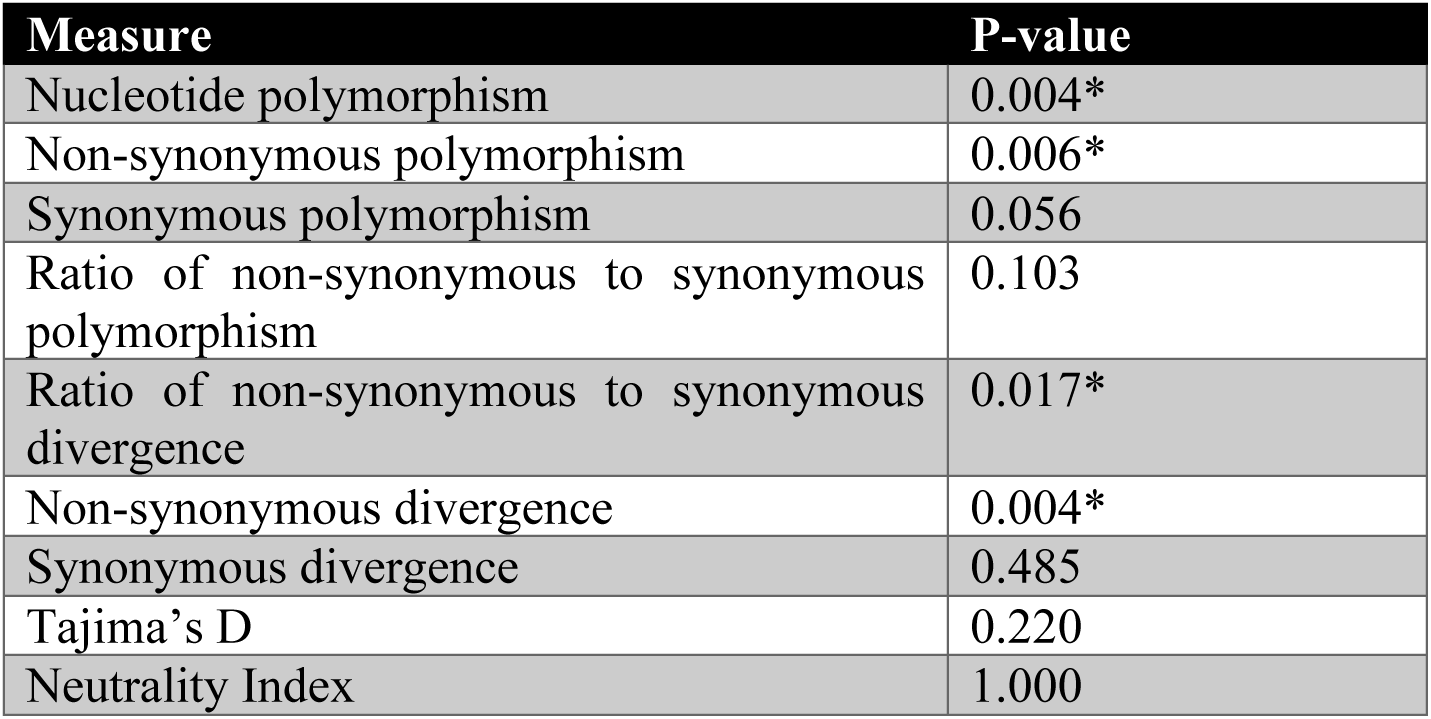

We find the same pattern as in the main analysis, with cooperative genes having a signature of higher non-synonymous polymorphism and divergence, without evidence for increased likelihood of positive or balancing selection.

This comparison is less well-controlled than the main analysis, but these results add further weight to our conclusion that signature of selection we observed in quorum sensing - controlled genes is a signature of kin selection.

### S3: Balancing selection

A logical explanation for cooperative genes to be more polymorphic than private genes would be that cooperative genes are more likely to be under balancing selection. This could occur if both cheats and cooperators are maintained in a population because the fitness advantage of cheats declines as they become more common (124, 125). It could also occur if multiple greenbeard recognition alleles are maintained (126).

To detect balancing selection from sequence alignment data, we can use several population genetic measures such as Tajima’s D and Fu & Li’s F* and D* that look at allele frequencies to determine if balancing selection is occurring. Tajima’s D looks at the distribution of allele frequencies, whereas Fu & Li’s measures look at singletons (rare variants found in only one strain). To interpret these measures, we use statistical tests to determine if each gene is significantly different from the neutral expectation. We then extract the list of genes with significant support, and test if they are overrepresented for social genes using binomial tests.

#### Tajima’s D

We use the beta-distribution test with alpha=0.025 from the Pegas package in R (127, 128) to identify which genes have evidence for balancing selection. The test could be significant either due to balancing selection and a lack of rare alleles (D>>0) or a recent selective sweep (D<<0), so we exclude significant genes where D<0.

We have 242 genes with significant evidence for balancing selection. Only one of those is a cooperative gene, so cooperative genes are not significant overrepresented in those under balancing selection (binomial test, p=0.380). That gene is the aminoglycoside resistance gene aadK.

#### Fu & Li’s D* and F*

We use the critical values from (129) for n=100 genes and alpha=0.025, which is 1.53 for D* and 1.73 for F*. The probability of D* or F* being greater than this critical value for by chance is 0.025. Although we have >100 genes, this is likely to be a good approximation as the critical value scales with the natural-log of n. Although we have >100 genes, this is likely to be a good approximation as the critical value scales with the natural-log of n.

We have 21 genes with significant evidence for balancing selection from Fu & Li’s D*. None of them are cooperative. We have 13 genes with significant evidence for balancing selection from Fu & Li’s D*. None of them are cooperative

Overall, these results support our conclusions that cooperative genes have higher polymorphism due to relaxed selection, rather than balancing selection.

### S4: Positive selection

Cooperative genes might show greater divergence than private genes due to being more likely to be under positive or directional selection.

We use several complementary measures to look for signatures of positive selection in our genes. First, we use the Mcdonald-Kreitman (MK) test, which compares the ratio of non-synonymous to synonymous divergence with the ratio of non-synonymous to synonymous polymorphism (130). If there is lots more non-synonymous divergence than non-synonymous polymorphism, then this could indicate positive selection. The logic is that with strong positive selection, advantageous mutations don’t spend much time as polymorphisms, and are mainly detected through divergence..

We have 15 genes which have evidence for significant positive selection from the MK test. None of them are cooperative.

Secondly, we use the neutrality index, which uses the same information as the MK test, but rather than just a binary significance test, it uses the full information to make a continuous variable (130).. This can be interpreted by comparing averages and distributions of different groups of genes. We log-transformed neutrality index to normalise, meaning that positive values indicate positive selection. There is no difference in neutrality index between cooperative and private genes, and both also don’t differ from the background set of genes (Kruskal-Wallis test, chi-squared=0.06, df=2, p=0.974) Supplementary Figure 4.1.

**Supplementary Figure 4.1:**
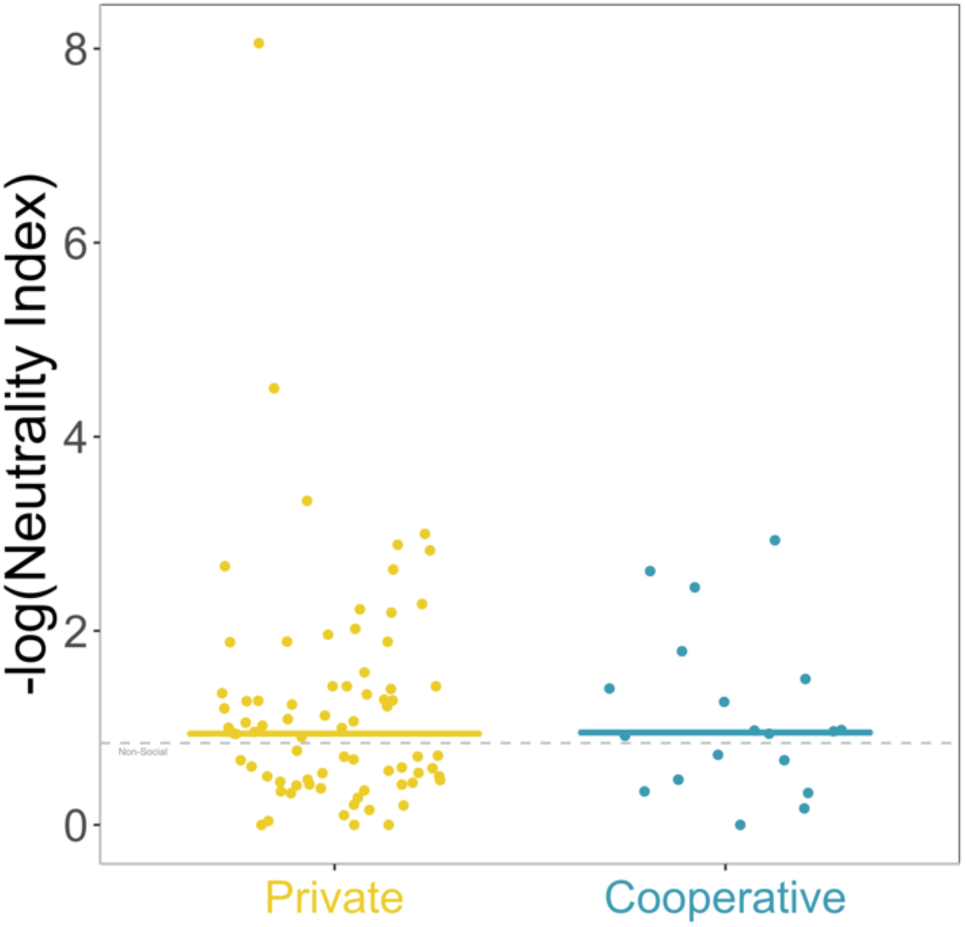
Log-transformed neutrality index of private and cooperative genes. The dashed line shows the median for background genes.

We also use the direction of selection statistic, which again uses the same information as the MK test to make a continuous variable, with positive values indicating positive selection. Whilst there is a slight trend for cooperative genes to have a higher direction of selection statistic than private genes, this is not significant (Kruskal-Wallis test, chi-squared=0.09, df=2, p=0.955) Supplementary Figure 4.2.

**Supplementary Figure 4.2:**
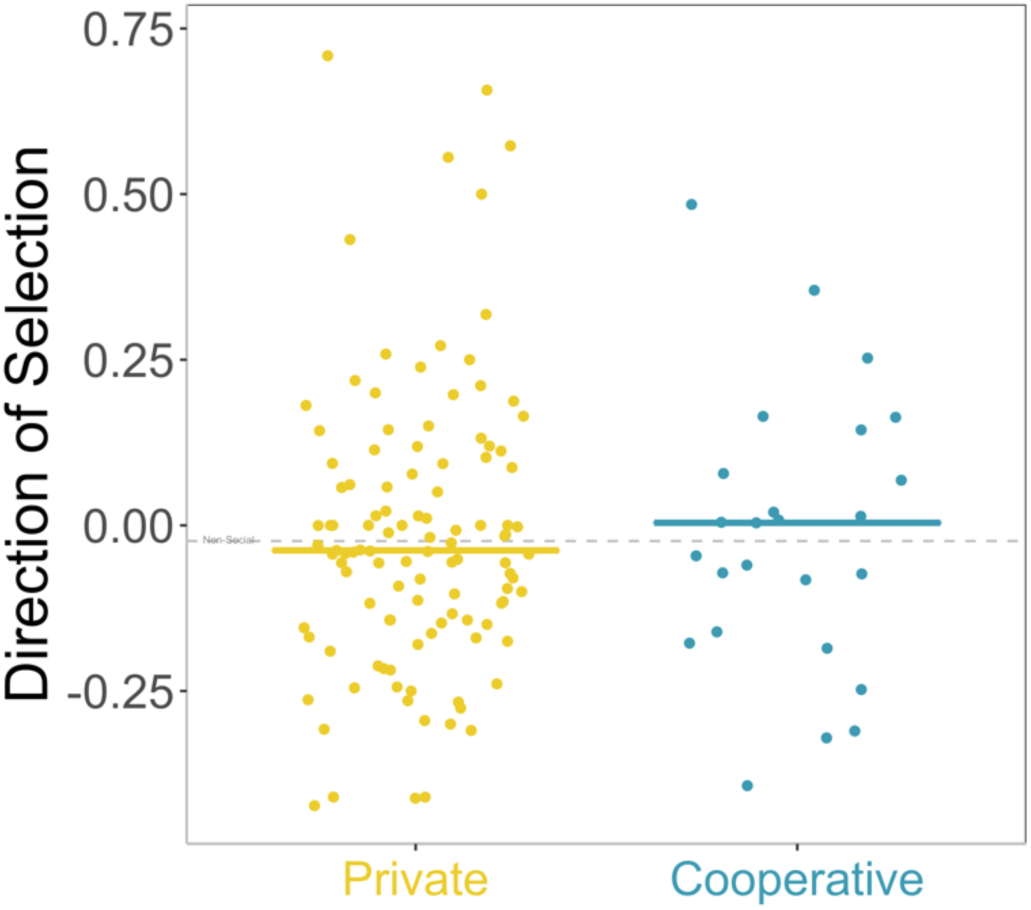
Direction of selection statistic for private and cooperative genes. The dashed line shows the median for background genes.

Overall, these findings support our conclusion that cooperative genes are not more divergent than private genes due to being more likely to be under positive selection.

### S5: Alternative explanations

#### Gene length

It is well-known that gene length can affect molecular population genetic parameters such as polymorphism. Even though we calculate all measures per site (considering gene length), we also check here that our results aren’t an artefact of any differences in gene length between cooperative and private genes.

Cooperative genes are on average 19% longer than private genes (965 base-pairs compared to 812).

We conduct a small analysis where we remove the smallest 25% of genes from our analysis, which is those genes which are <450 base-pairs long.

Cooperative genes are still significantly more polymorphic and divergent than private genes using this reduced dataset (Kruskal-Wallis test, chi-squared=8.16, df=2, p=0.017. Dunn Test p=0.01)

#### Horizontal gene transfer / pangenome

We focused our analysis on chromosomal genes, because frequent horizontal gene transfer can make drawing conclusion from molecular population genetic parameters more challenging. We check if cooperative genes are more likely to be horizontally transferred by checking if they are overrepresented in the accessory genome compared to the core genome.

*B. subtilis* has an open pangenome, meaning that each additional strain sequenced adds many new genes. This is usually indicative of a species living in multiple or variable environments, which we know is true for this species. *B. subtilis* is also naturally competent, meaning they can take-up DNA from the environment, and it is thought that the open pangenome occurs because rare genes are acquired in this way from closely related species (94).

We use the panX database (pangenome.org) to assign genes as core or accessory genome based on 80 *B. subtilis* genomes. If we count core genes as those present in 90% of genomes, then 23.2% of the genes in our reference strain are in the *B. subtilis* accessory genome, and 76.8% are in the core genome.

10 out of 53 cooperative genes are in the accessory genome, which is 18.9%. Cooperative genes are therefore not overrepresented in the accessory genome (binomal test p=0.622).

We conclude that cooperative genes are not more likely to be transferred horizontally, which matches with previous results (78)..

### Power analysis

We conducted a power analysis to see what would happen if we had fewer strains in our population genetic analysis.

We took advantage of the vcf-tools software, which allowed us to randomly remove strains from our analysis. This enabled us to conduct a basic analysis on polymorphism for a groups of N strains (N = 4,6,8,10,12,14,16,18,20,22,24,26,28), with 22 iterations of each number of strains

Within any analysis, we focussed on the 3570 genes which are present in all strains, and calculated mean and median polymorphism (average pairwise polymorphism, relative to gene length) (Supplementary Figure S5.1).

**Supplementary Figure S5.1:**
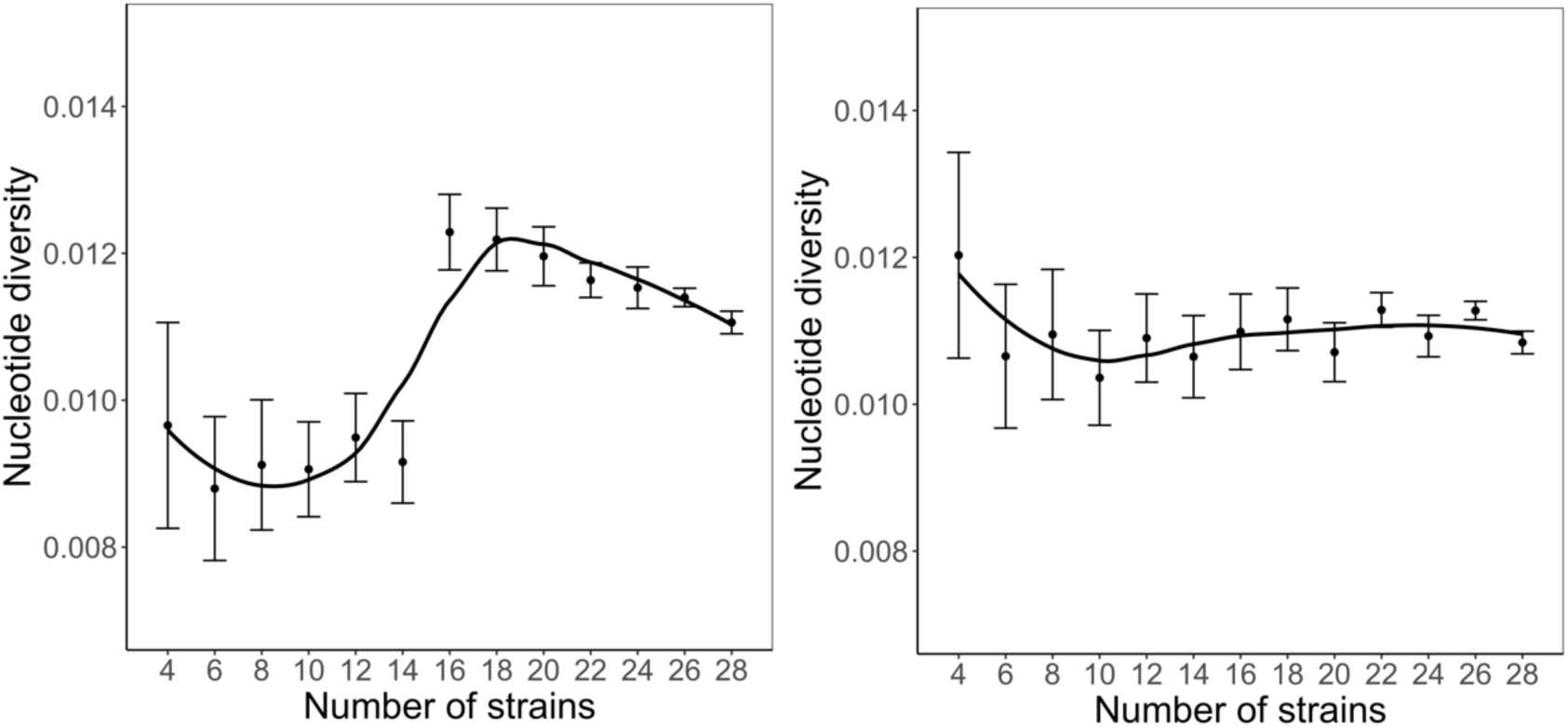
(A) Median polymorphism, (B) Mean polymorphism as the number of strains uses in the analysis varies. The line is a loess regression fit

The graph on the left shows median polymorphism, which shows a threshold effect once we get to 16 strains, and also much smaller error bars. This shows that a smaller number of strains will miss a lot of the diversity between strains.

The graph on the right shows mean polymorphism. The main pattern is that the standard error in mean polymorphism declines substantially as the number of strains increase. This means that the likelihood of getting a good estimate of the population mean increases as the number of strain increases, which makes logical sense.

The fact that mean polymorphism doesn’t change much, but median does, implies that the distribution has changed. The overall conclusion is that the number of strains we have included is likely appropriate to capture the true variation in the population

### S6: Division of labour

We conducted an analysis to see if the division of labour in the production of public goods in *B. subtilis* could explain the signatures of selection that we observe. For this, we look at the subset of 40 genes which are controlled by Spo0A, which is active only in a subset of cells (63). Depending on whether Spo0A is on or off, the expression of extracellular polysaccharides and amyloid protein fibres is either repressed or not (32, 61).

Spo0A controlled genes have lower median polymorphism than other quorum sensing - controlled genes, whether measured as overall polymorphism (0.0045 vs. 0.0101), non-synonymous polymorphism (0.0015 vs. 0.0037), or synonymous polymorphism (0.016 vs. 0.021).

Spo0A controlled genes also have lower non-synonymous divergence (0.013 vs. 0.023) and lower ratio between non-synonymous and synonymous divergence (0.049 vs. 0.097), although they have slightly higher synonymous divergence (0.286 vs. 0.266).

The direction of selection statistic is very similar between the two groups (-0.044 vs. -0.037).

This pattern is the opposite to what we would expect if the division of labour was responsible for the signature of selection, rather than sociality *per se*. If the lower conditional expression was having a large effect, then we would expect these genes to have higher polymorphism than the background set of quorum sensing-controlled genes.

Further, EPS and TasA genes stand out within this class of genes as having high polymorphism, implying that the social effect is important in causing the signature of selection that we observe.

### S7: Competence

The reference strain NCIB 3610 has a plasmid-encoded gene ComI, which interferes with the competence machinery (Konkol 2013). We know that there is variation in natural competence in our strains, as they were all screened for competency by (35). 18 out of the 31 strains we used are genetically competent.

Some of the competence genes (comGF, comGE, comGG) have extremely high polymorphism. Here, we conduct a small analysis where we restrict our analysis to only competent strains, to see if this polymorphism is caused by disuse in non-competent strains. We find that the competence operon comG is still amongst the most polymorphic even when we are only looking at competent strains.

Furthermore, cooperative genes are still significantly more polymorphic than private genes in the competent strains (Kruskal-Wallis test, chi-squared=14.7, df=2, p<0.0001).

Overall, we conclude that variation in natural competence isn’t responsible for the difference between cooperative and private genes that we observe.

### S8: Estimation of relatedness

According to theory, the degree to which selection is relaxed in cooperative genes relative to private genes is inversely proportional to relatedness. This result emerges from a simple population genetics model, which shows that a slightly deleterious allele with a cooperative effect on fitness will reach equilibrium frequency inversely proportional to the relationship between the actor and recipient (r) (19). If we assume weak selection and a large population (and ignore higher-order terms), we can directly map this prediction to relative levels of nucleotide polymorphism between alleles with cooperative and private effects on fitness.

As we noted in our previous work on *P. aeruginosa,* we have to make further assumptions that our set of cooperative genes experience the same average strength of selection and distribution of fitness effects as private genes, but this approach has the advantage of many experimental attempts to estimate relatedness for social interactions in that we don’t need to know the scale at which interactions take place, or the relative weighting of environments in which traits are more or less social.

Cooperative genes have a median polymorphism of 0.01078, and private genes have median polymorphism of 0.00918. This leads to a calculation of relatedness as r=0.77.

We note that this measure might vary depending on how you define the population, which is tricky in bacteria due to horizontal gene transfer and other complications (131). In *B. subtilis*, for example, we know that strains from the same plant root or gram of soil can vary in their production of public goods, and don’t always group together phylogenies (132–134).

### S9: Correlation in gene expression for secondary comparisons

Here, we use the dataset from Futo *et al*. 2020 to test if the genes we used in the secondary comparisons also tend to be co-expressed. For each set of genes we first calculate the mean pairwise correlation between all possible pairs of genes. We then use a bootstrap approach of randomly sampling 10,000 other gene sets of the same size, to see if the gene set has higher correlated expression than expected by chance.

For all gene sets, we find that the correlation is greater than >95% of randomly sampled gene-sets (Supplementary Table S9.1)

**Supplementary Table S9.1:** Results from pairwise correlation in gene expression of a gene set, and bootstrap iterations of randomly sampled gene sets of the same size

| Gene set | Correlation for gene set | Mean correlation for 10,000 iterations | Percentile of gene set |
| --- | --- | --- | --- |
| Iron scavenging | 0.880 | 0.329 | 100 |
| Antibiotic resistance | 0.429 | 0.306 | 95.7 |
| Proteases | 0.397 | 0.293 | 95.3 |
| Toxins | 0.534 | 0.305 | 99.8 |
| Antimicrobials | 0.640 | 0.294 | 100 |

### S10: Positive and balancing selection in other cooperative traits

We checked whether the other cooperative traits that we examined differ in signatures of positive and balancing selection, as this could cloud our conclusion on the nature of selection. We use Tajima’s D to detect balancing selection, and neutrality index to detect positive selection.

Cooperative genes don’t differ in balancing selection, whether we use all genes from the six comparisons (ANOVA *F*_2,121_ = 7.50, *p* < 0.001; Games Howell Test p=0.39), just the five secondary comparisons (ANOVA *F*_2,44_ = 4.72, *p* = 0.14; Games Howell Test p=0.33), or consider each gene as a data point (Wilcoxon signed-rank test V=7, p=0.563).

Cooperative genes also don’t differ in positive selection, whether we use all genes from the six comparisons (Kruskal-Wallis *X*^2^(2) = 2.11, *p* = 0.35, Dunn Test p = 0.70), just the five secondary comparisons (Kruskal-Wallis *X*^2^(2) = 5.37, *p* = 0.07, Dunn Test p = 0.98), or consider each gene as a data point (Wilcoxon signed-rank test V=7, p=0.563).

Overall, we can conclude that the signature of selection we see across the five other cooperative traits are most consistent with kin selection causing the effective relaxation of selection

### S11: Correlations in gene expression of QS-controlled genes

To test the robustness of our result that quorum sensing-controlled genes tend to be expressed at the same time, we repeated our analysis using the dataset from Pisithkul (37).

Whilst Futo *et al*. measured gene expression in a solid-air interface biofilm over two months (N=11 timepoints), Pisithkul *et al*. used a liquid-air interface, and measured gene expression over 24 hours (N=7 timepoints), representing the initial stages of biofilm growth.

Pisithkul *et al*. provided their data in the form of reads per kilobase of transcript per million mapped reads. For each gene, we normalised the data with the following steps;

(1) We calculated median expression across the four replicates of each timepoint
(2) We then divided each measure by the median expression at timepoint one (eight hours). This allows for better comparison between genes
(3) We then used a log2 transformation to normalize the resulting relative expression measures

The average correlation for the N=160 quorum sensing controlled genes that we were able to match to the data was 0.411. The average correlation for N=10,000 randomly sampled gene sets of the same size (N=160) was 0.385. Quorum sensing-controlled genes have a higher correlation than 98.5% of the randomly sampled gene sets.

## Supplementary Figures

**Supplementary Figure 1:**
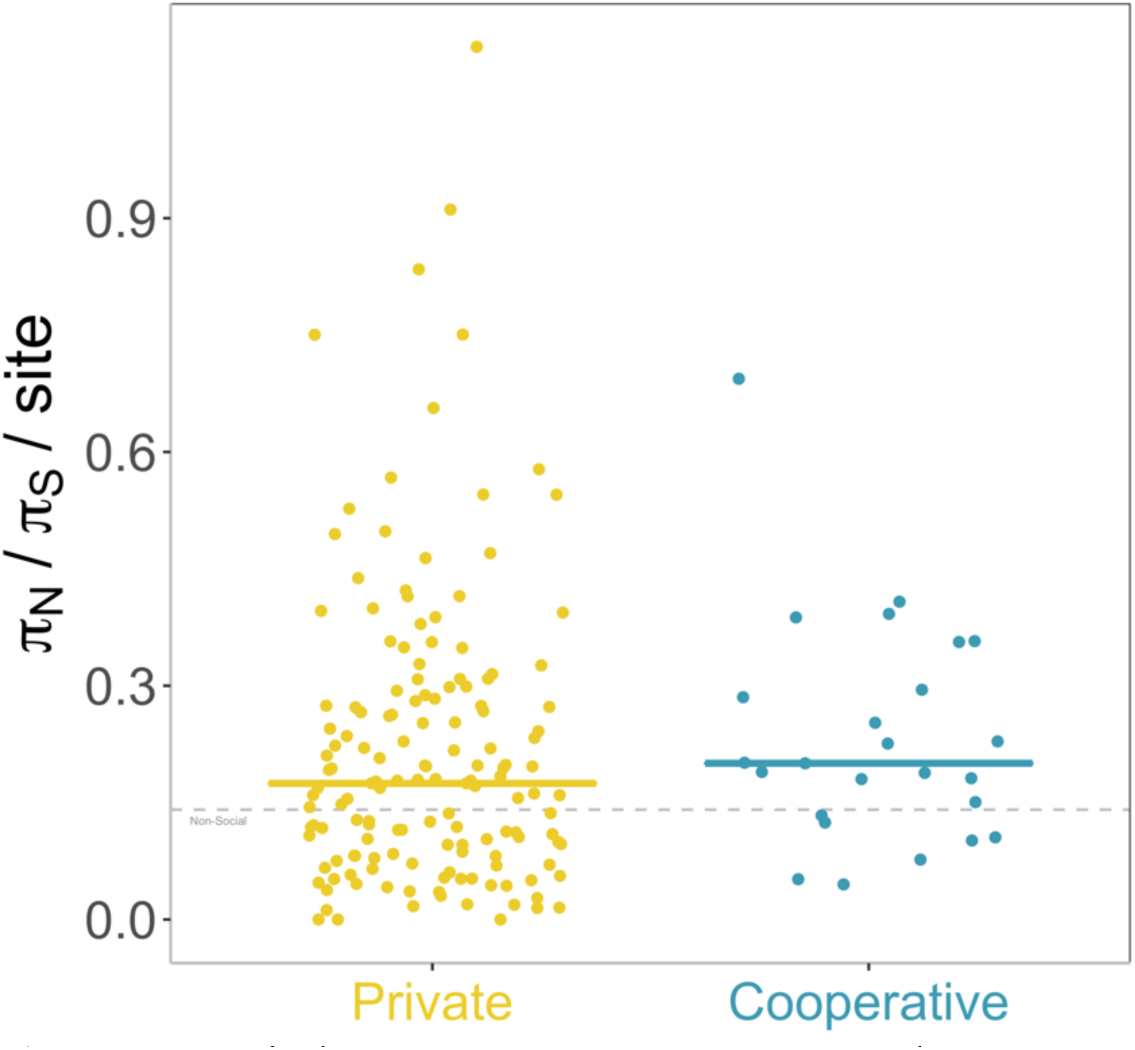
Ratio between non-synonymous and synonymous nucleotide diversity per site for private (yellow) and cooperative (blue) genes controlled by quorum sensing. Each point is a gene, and the horizontal line shows the median for each group. The grey line shows the median for background private genes across the genome.

**Supplementary Figure 2:**
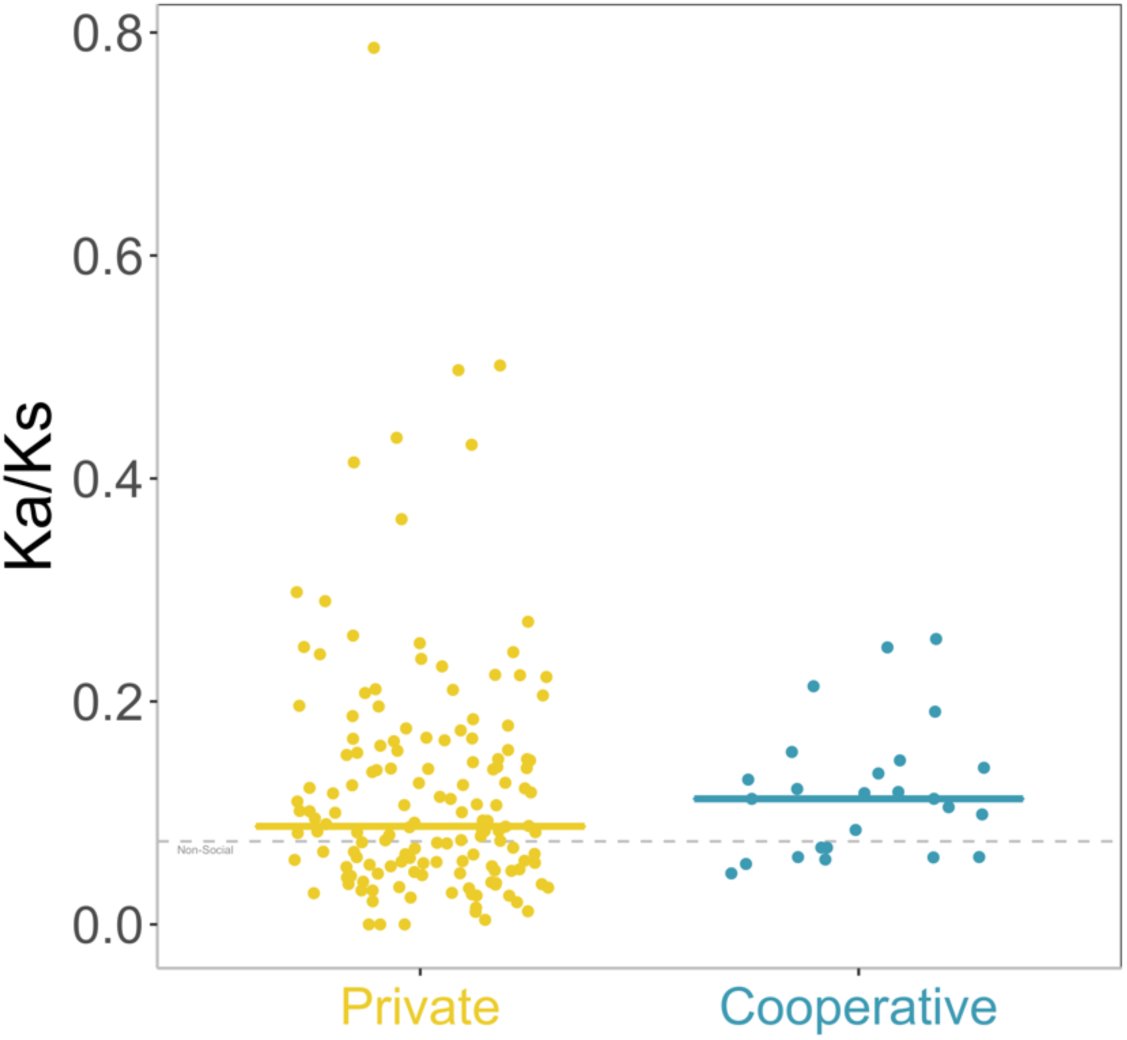
Ratio between non-synonymous and synonymous divergence for private (yellow) and cooperative (blue) genes controlled by quorum sensing. Divergence is measured by rates of protein evolution, e.g. number of synonymous substitutions per synonymous site for panel B. Each point is a gene, and the horizontal line shows the median for each group. The grey line shows the median for background private genes across the genome.

**Supplementary Figure 3:**
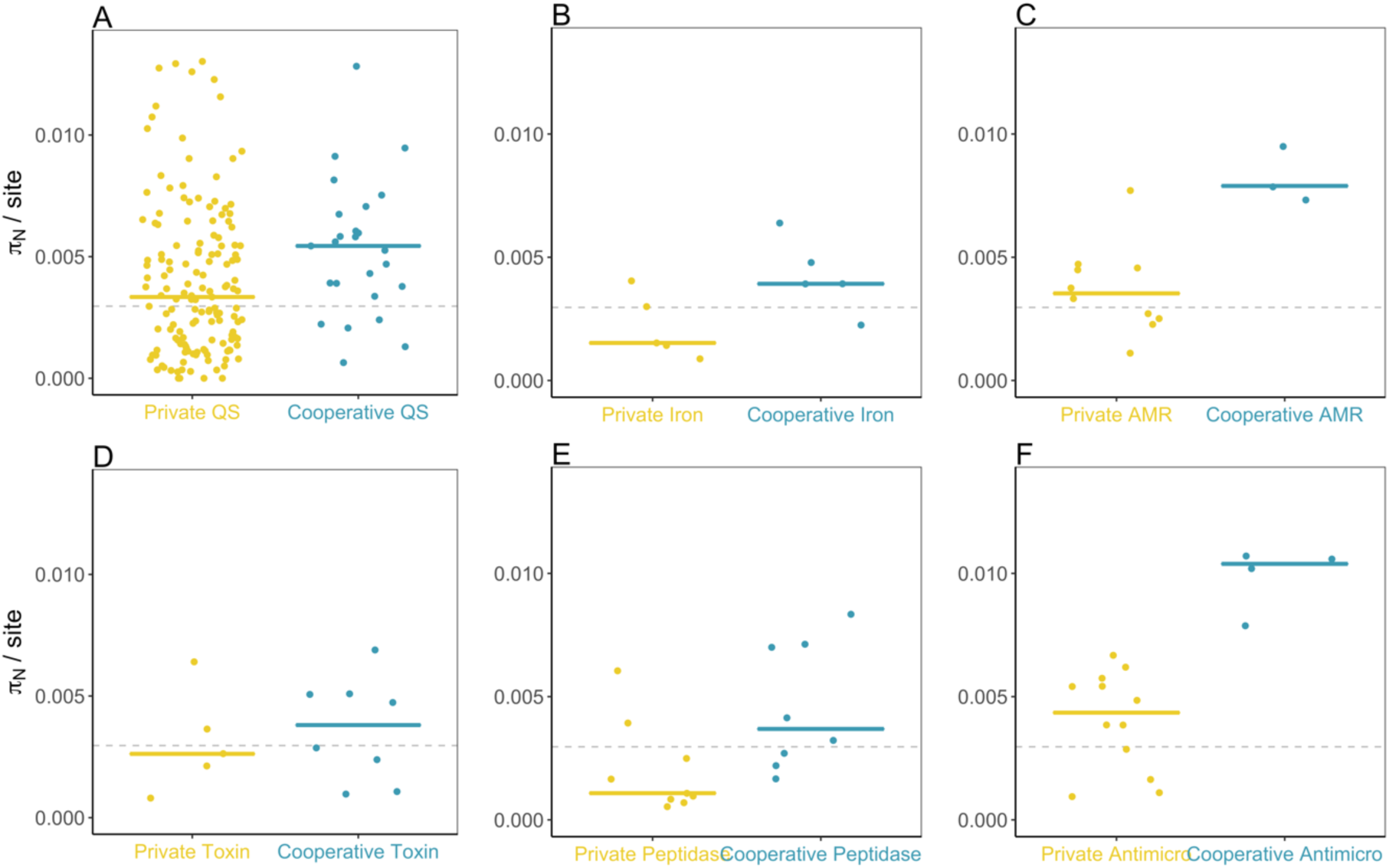
Private (yellow) vs. cooperative (blue) non-synonymous polymorphism in genes for six traits. Panel A shows the quorum sensing -controlled genes used in the main analysis. Panels B-F show the secondary comparisons

**Supplementary Figure 4:**
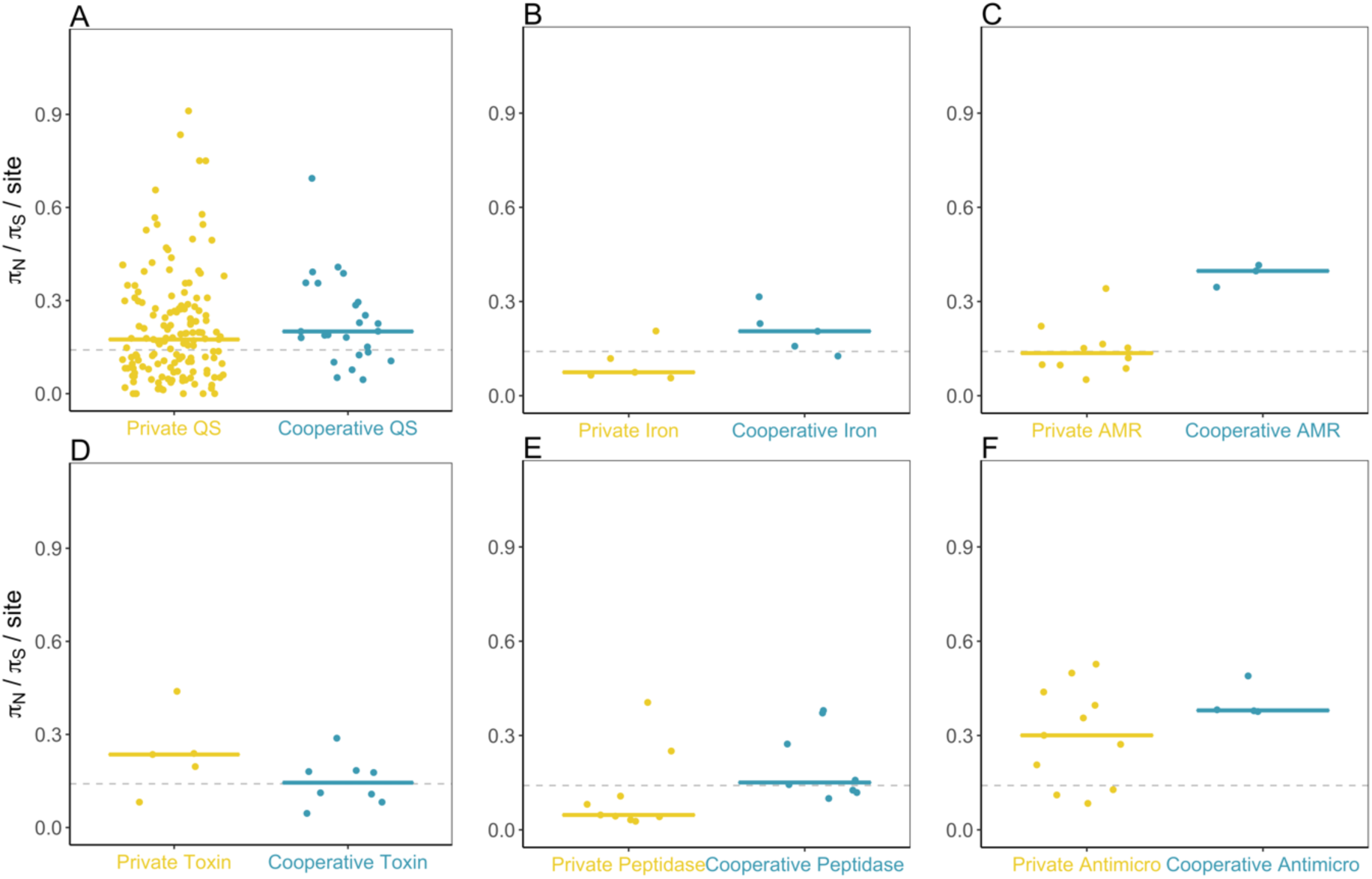
Private (yellow) vs. cooperative (blue) ratio between non-synonymous and synonymous polymorphism in genes for six traits. Panel A shows the quorum sensing-controlled genes used in the main analysis. Panels B-F show the secondary comparisons

**Supplementary Figure 5:**
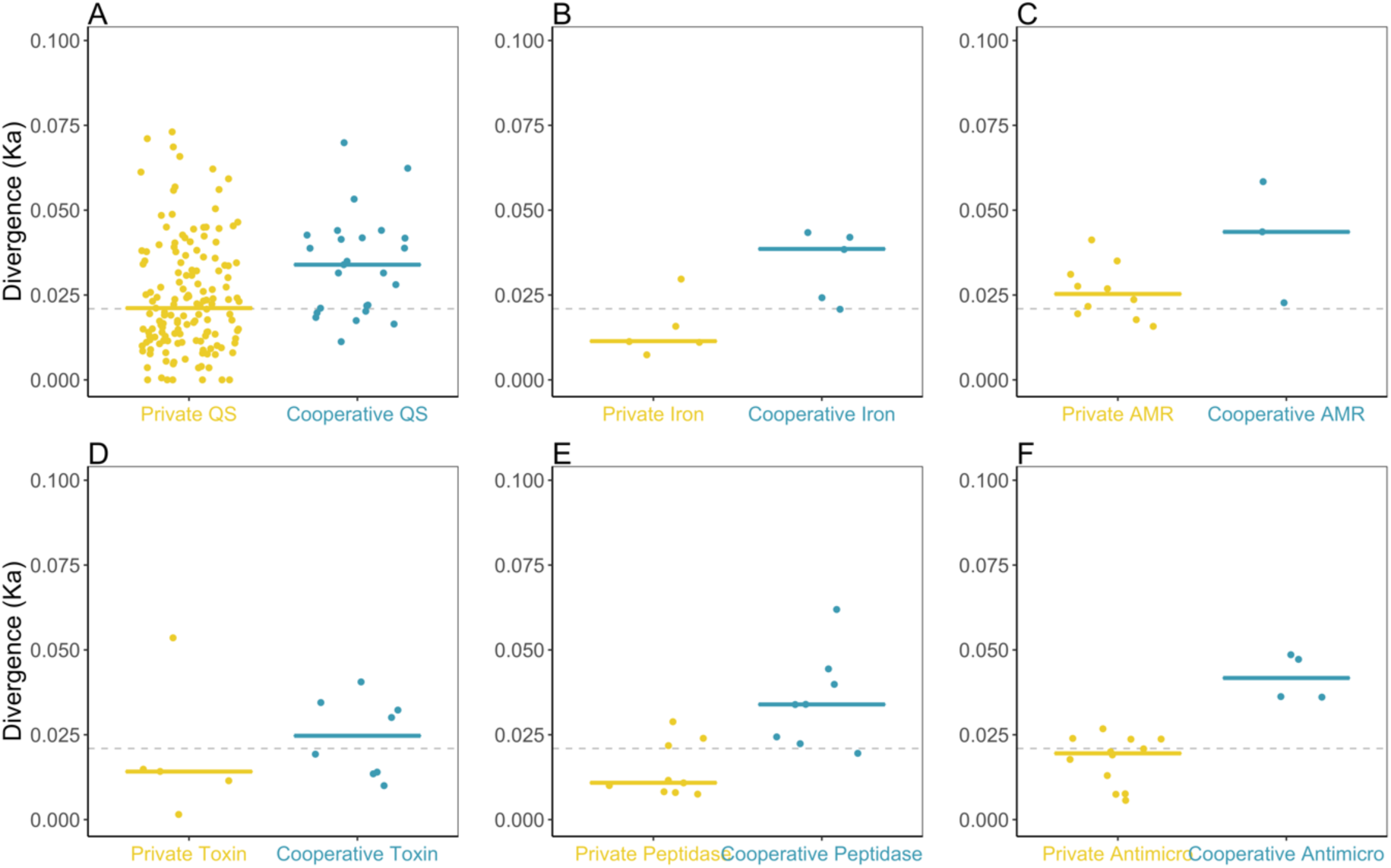
Private (yellow) vs. cooperative (blue) non-synonymous divergence in genes for six traits. Panel A shows the quorum sensing -controlled genes used in the main analysis. Panels B-F show the secondary comparisons

**Supplementary Figure 6:**
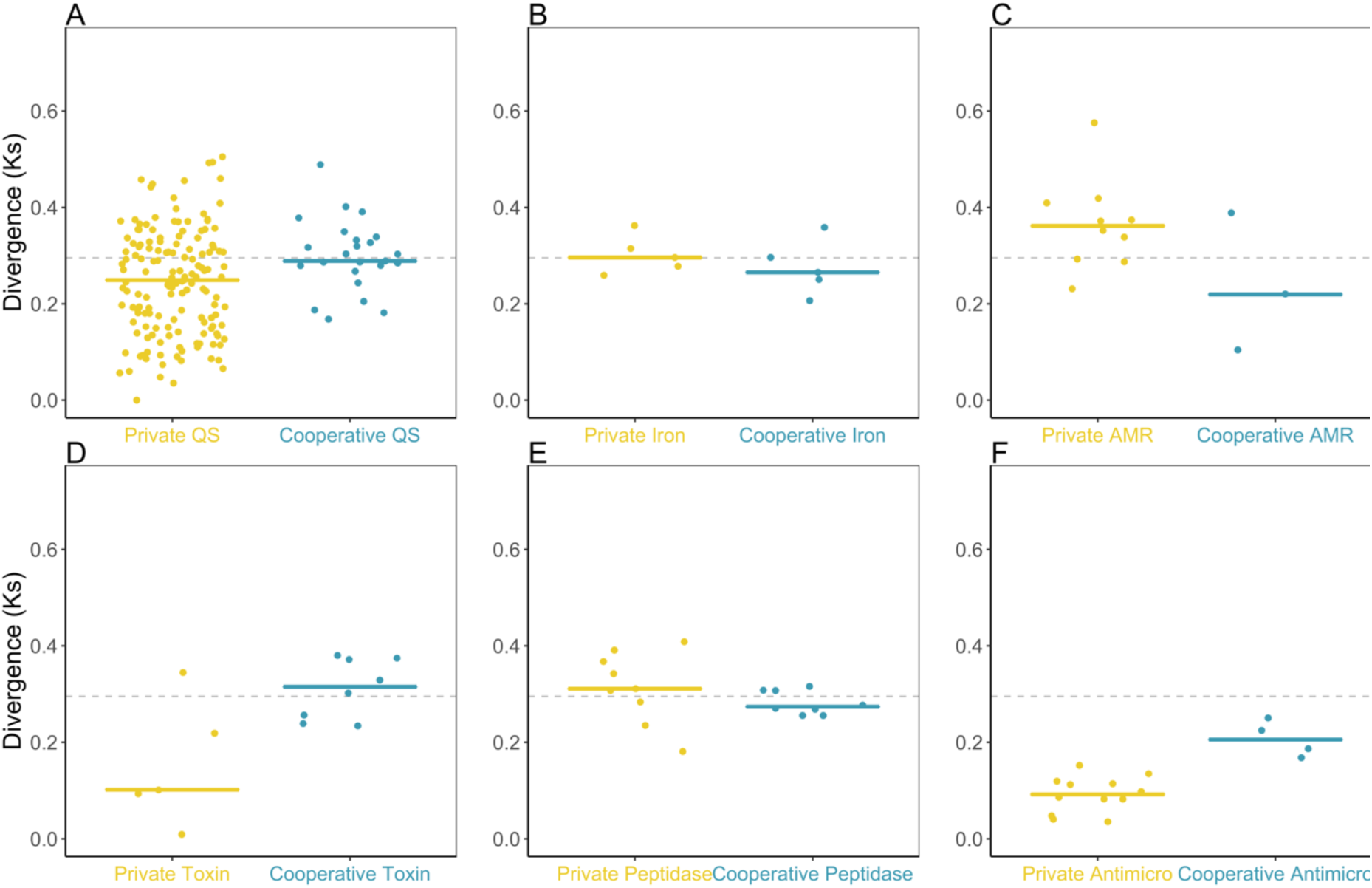
Private (yellow) vs. cooperative (blue) synonymous divergence in genes for six traits. Panel A shows the quorum sensing -controlled genes used in the main analysis. Panels B-F show the secondary comparisons

**Supplementary Figure 7:**
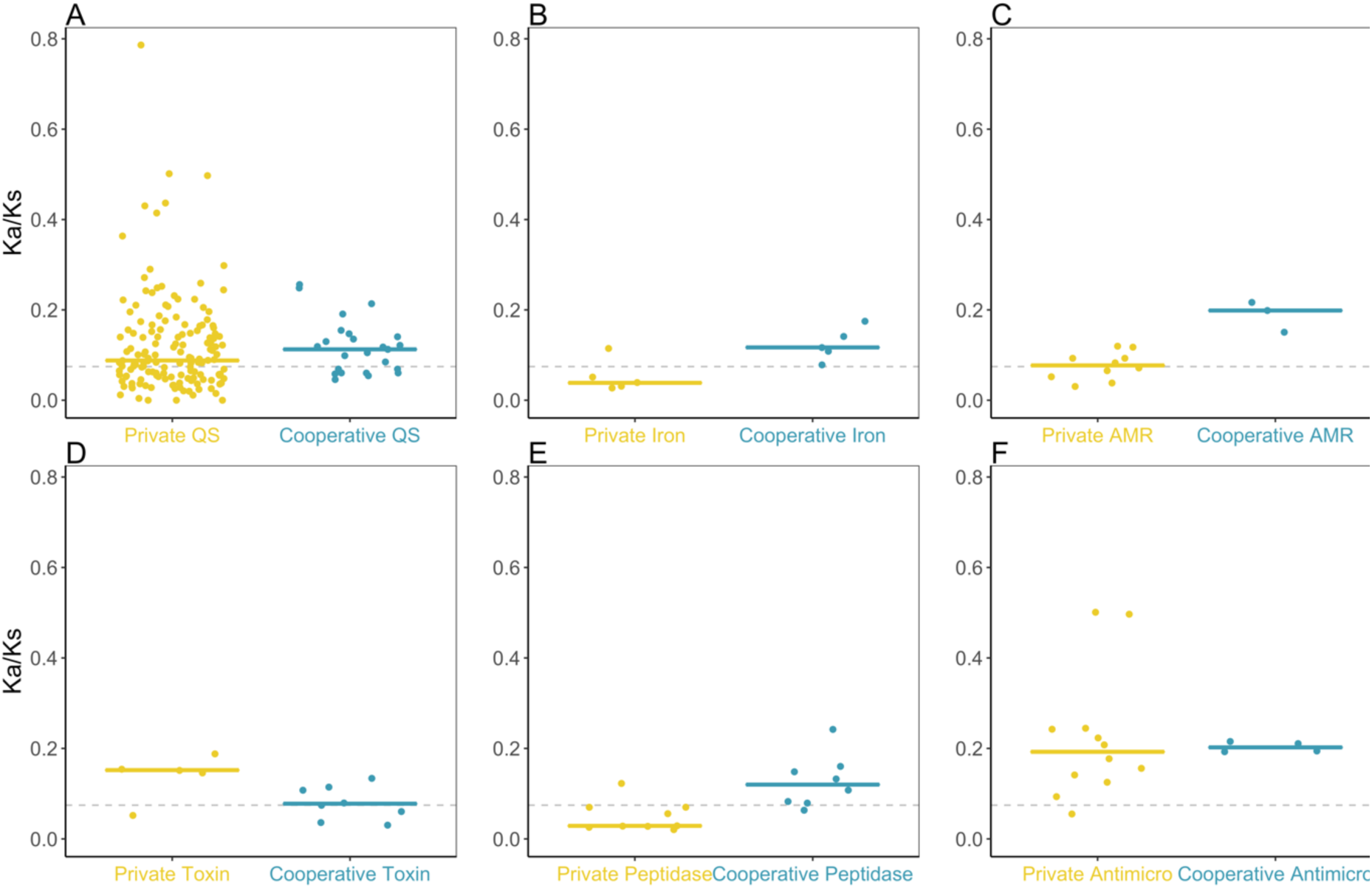
Private (yellow) vs. cooperative (blue) ratio between non-synonymous and synonymous divergence in genes for six traits. Panel A shows the quorum sensing-controlled genes used in the main analysis. Panels B-F show the secondary comparisons

## Supplementary Tables

**Supplementary Table 1:** List of strains used

| Strain ID | Location | Assembly |
| --- | --- | --- |
| NRS6096 | Tayport (UK) garden soil | GCA_905311035.1 |
| NRS6099 | Tayport (UK) garden soil | GCA_905310985.1 |
| NRS6103 | Tayport (UK) community garden soil | GCA_905310995.1 |
| NRS6105 | Tayport (UK) community garden soil | GCA_905311425.1 |
| NRS6107 | Tayport (UK) garden soil | GCA_905311405.1 |
| NRS6108 | Tayport (UK) garden soil | GCA_905311395.1 |
| NRS6110 | Tayport (UK) vegetable plot | GCA_905311415.1 |
| NRS6111 | Tayport (UK) vegetable plot | GCA_905311375.1 |
| NRS6116 | Tayport (UK) garden soil | GCA_905311385.1 |
| NRS6118 | Tayport (UK) potato patch | GCA_905311435.1 |
| NRS6120 | Tayport (UK) potato patch | GCA_905312035.1 |
| NRS6121 | Tayport (UK) shrub bed | GCA_905315035.1 |
| NRS6127 | Tayport (UK) garden soil | GCA_905315045.1 |
| NRS6128 | Tayport (UK) garden soil | GCA_905315055.1 |
| NRS6131 | Tayport (UK) worm bin | GCA_905315025.1 |
| NRS6134 | Tayport (UK) vegetable patch | GCA_905315395.1 |
| NRS6137 | Tayport (UK) community garden soil | GCA_905315385.1 |
| NRS6141 | Tayport (UK) garden soil | GCA_905315375.1 |
| NRS6145 | Tayport (UK) garden soil | GCA_905315685.1 |
| NRS6148 | Tayport (UK) garden soil | GCA_905315695.1 |
| NRS6153 | Tayport (UK) garden soil | GCA_905315705.1 |
| NRS6160 | Tayport (UK) garden soil | GCA_905315715.1 |
| NRS6167 | Tayport (UK) soil from planter | GCA_905316385.1 |
| NRS6181 | Tayport (UK) garden soil | GCA_905318255.1 |
| NRS6183 | Lochee (UK) garden soil | GCA_905319155.1 |
| NRS6186 | Newport (UK) garden soil | GCA_905319135.1 |
| NRS6190 | Tayport (UK) garden soil | GCA_905319565.1 |
| NRS6194 | Tayport (UK) garden soil | GCA_905319555.1 |
| NRS6202 | Kirriemuir (UK) garden soil | GCA_905319535.1 |
| NRS6205 | Kirriemuir (UK) garden soil | GCA_905319825.1 |
| NRS6206 | Kirriemuir (UK) garden soil | GCA_905319815.1 |

**Supplementary Table 2:** List of social genes

| Gene | Function | Reference |
| --- | --- | --- |
| **bslA** | biofilm-surface layer protein BslA | (33, 34) |
| **bslB** | biofilm-surface layer protein BslB | (135) |
| **epsA** | hypothetical protein | (33, 34) |
| **epsB** | protein tyrosine kinase EpsB | (33, 34) |
| **epsC** | polysaccharide biosynthesis protein | (33, 34) |
| **epsD** | glycosyltransferase family 4 protein | (33, 34) |
| **epsE** | glycosyltransferase EpsE | (33, 34) |
| **epsF** | glycosyltransferase family 1 protein | (33, 34) |
| **epsG** | biofilm exopolysaccharide biosynthesis protein EpsG | (33, 34) |
| **epsH** | glycosyltransferase | (33, 34) |
| **epsI** | polysaccharide pyruvyl transferase family protein | (33, 34) |
| **epsJ** | lipoprotein | (33, 34) |
| **epsK** | cyclic-di-GMP receptor EpsK | (33, 34) |
| **epsL** | sugar transferase | (33, 34) |
| **epsM** | acetyltransferase | (33, 34) |
| **epsN** | aminotransferase class I/II-fold pyridoxal phosphate-dependent enzyme | (33, 34) |
| **epsO** | polysaccharide pyruvyl transferase family protein | (33, 34) |
| **sipW** | signal peptidase I | (33, 34) |
| **srfAA** | surfactin non-ribosomal peptide synthetase SrfAA | (33, 34) |
| **srfAB** | surfactin non-ribosomal peptide synthetase SrfAB | (33, 34) |
| **srfAC** | surfactin non-ribosomal peptide synthetase SrfAC | (33, 34) |
| **srfAD** | surfactin biosynthesis thioesterase SrfAD | (33, 34) |
| **srfT** | MFS transporter | (33, 34) |
| **tapA** | amyloid fiber anchoring/assembly protein TapA | (33, 34) |
| **tasA** | biofilm matrix protein TasA | (33, 34) |

**Supplementary Table 3:** Cooperative and private genes for iron-scavenging via bacillibactin

| Gene ID | Name | Function | Sociality | Reference |
| --- | --- | --- | --- | --- |
| **IRON SCAVENGING** |  |  |  |  |
| **17270** | dhbF | Bacillibactin biosynthesis | Cooperative | (65, 66) |
| **17275** | dhbB | Bacillibactin biosynthesis | Cooperative | (65, 66) |
| **17280** | dhbE | Bacillibactin biosynthesis | Cooperative | (65, 66) |
| **17285** | dhbC | Bacillibactin biosynthesis | Cooperative | (65, 66) |
| **17290** | dhbA | Bacillibactin biosynthesis | Cooperative | (65, 66) |
| **1040** | feuC | Membrane permease | Private | (65, 66) |
| **1045** | feuB | Membrane permease | Private | (65, 66) |
| **1050** | feuA | Periplasmic binding protein | Private | (65, 66) |
| **17785** | yusV [feuV] | ABC transporter | Private | (65, 66) |
| **17295** | besA | Ferri-bacillibactin esterase | Private | (65, 66) |

**Supplementary Table 4:**
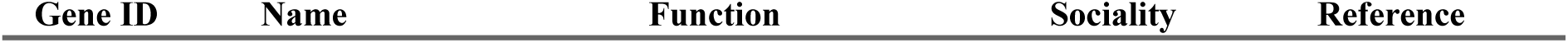
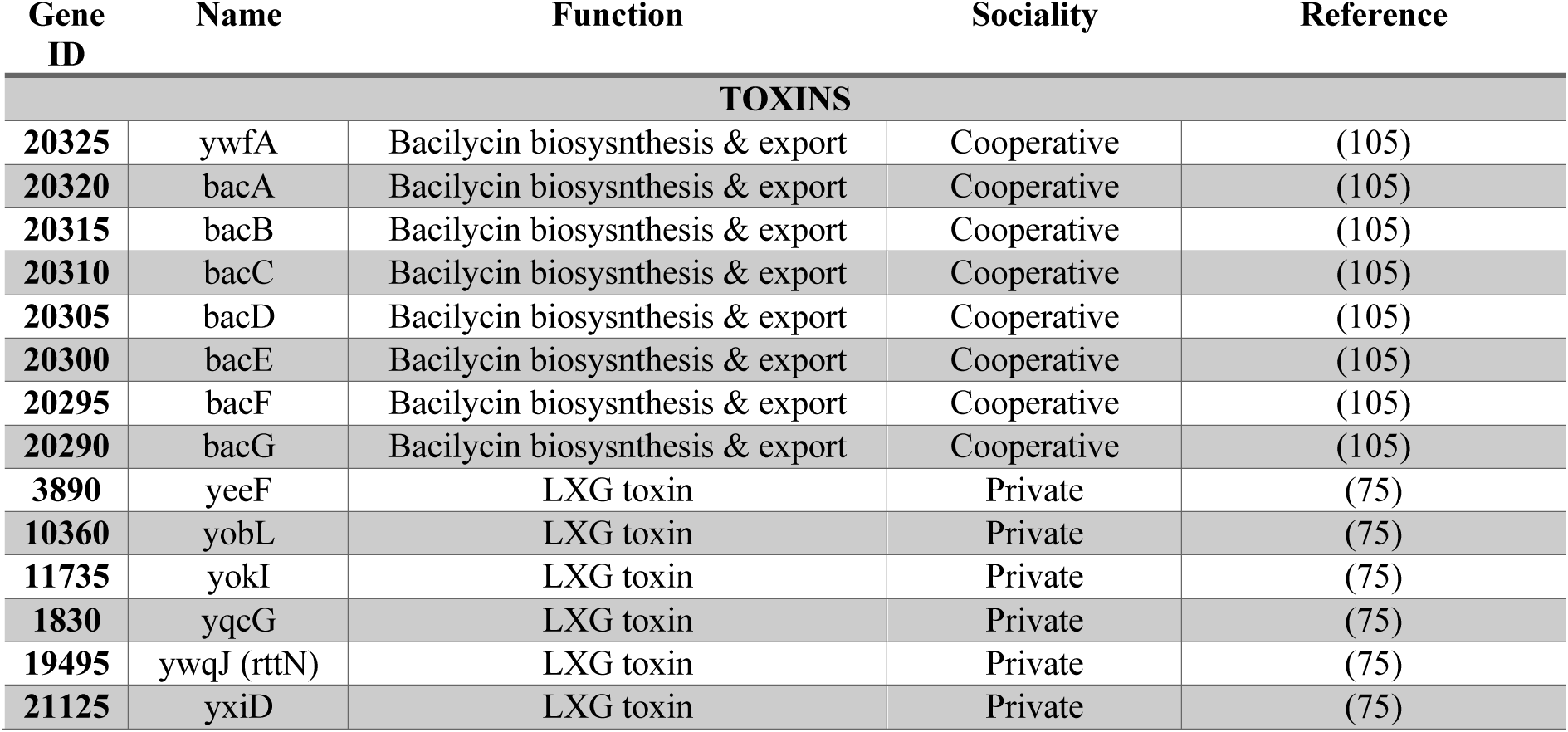
Cooperative and private genes for antibiotic resistance

**Supplementary Table 5:**
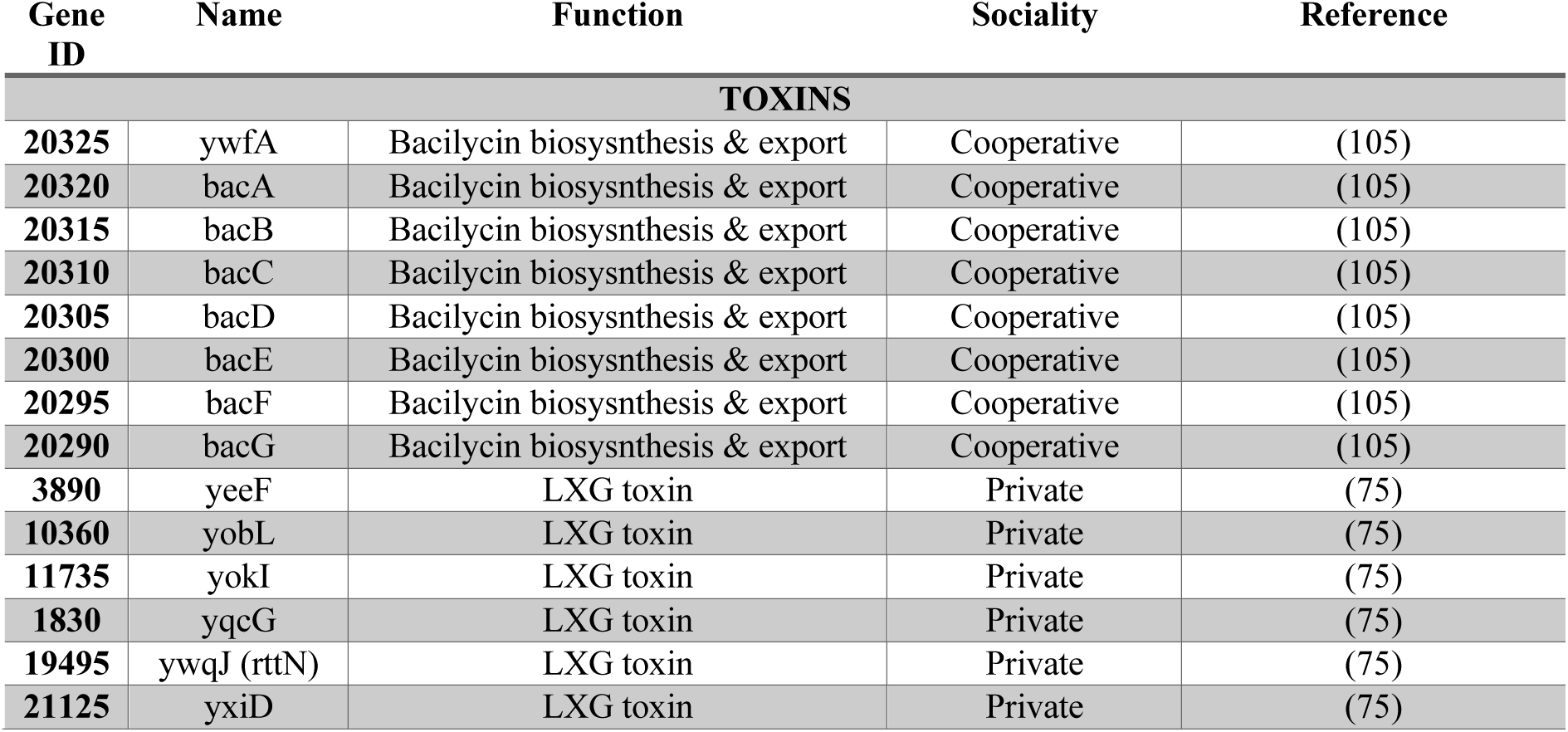
Cooperative and private genes toxin genes

**Supplementary Table 6:** Cooperative and private genes protease genes

| Gene ID | Name | Function | Sociality | Reference |
| --- | --- | --- | --- | --- |
| PROTEASES |  |  |  |  |
| 5755 | aprE | Extracellular protease | Cooperative | (70) |
| 8405 | Bpr | Extracellular protease | Cooperative | (70) |
| 20660 | Epr | Extracellular protease | Cooperative | (70) |
| 1375 | Mpr | Extracellular protease | Cooperative | (70) |
| 6155 | nprB | Extracellular protease | Cooperative | (70) |
| 8100 | nprE | Extracellular protease | Cooperative | (70) |
| 20500 | Upr | Extracellular protease | Cooperative | (70) |
| 5990 | wprA | Extracellular protease | Cooperative | (70) |
| 9385 | aprX | Intracellular protease | Private | (70) |
| 7560 | clpE | Intracellular protease | Private | (70) |
| 18630 | clpP | Intracellular protease | Private | (70) |
| **8830** | clpQ | Intracellular protease | Private | (70) |
| **13140** | ispA | Intracellular protease | Private | (70) |
| **15210** | lonA | Intracellular protease | Private | (70) |
| **15215** | lonB | Intracellular protease | Private | (70) |
| **9115** | mlpA | Intracellular protease | Private | (70) |
| **12000** | ypwA | Intracellular protease | Private | (70) |

**Supplementary Table 7:**
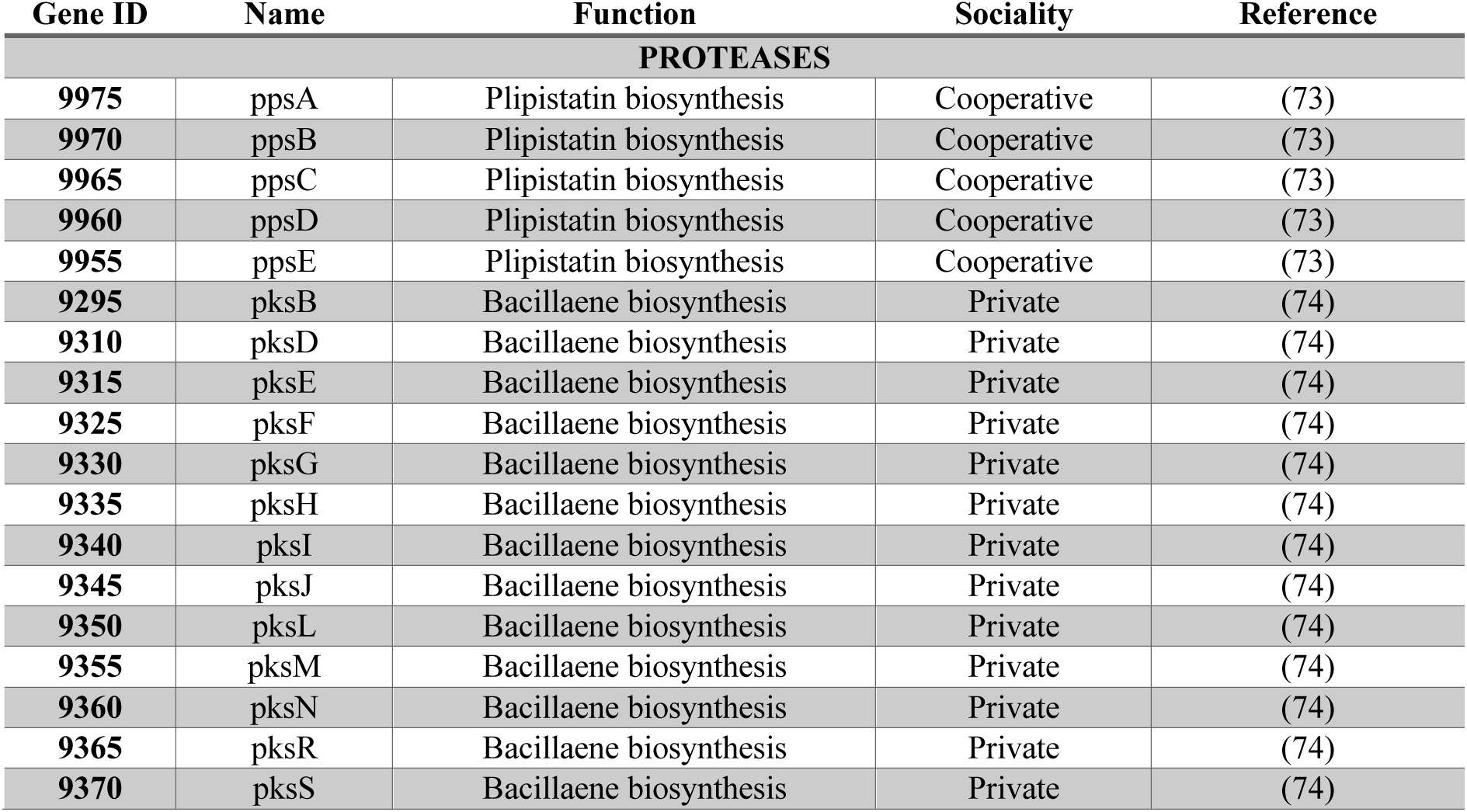
Cooperative and private genes for the production of antimicrobials

| Gene ID | Name | Function | Sociality | Reference |
| --- | --- | --- | --- | --- |
| **PROTEASES** |  |  |  |  |
| **9975** | ppsA | Plipistatin biosynthesis | Cooperative | (73) |
| **9970** | ppsB | Plipistatin biosynthesis | Cooperative | (73) |
| **9965** | ppsC | Plipistatin biosynthesis | Cooperative | (73) |
| **9960** | ppsD | Plipistatin biosynthesis | Cooperative | (73) |
| **9955** | ppsE | Plipistatin biosynthesis | Cooperative | (73) |
| **9295** | pksB | Bacillaene biosynthesis | Private | (74) |
| **9310** | pksD | Bacillaene biosynthesis | Private | (74) |
| **9315** | pksE | Bacillaene biosynthesis | Private | (74) |
| **9325** | pksF | Bacillaene biosynthesis | Private | (74) |
| **9330** | pksG | Bacillaene biosynthesis | Private | (74) |
| **9335** | pksH | Bacillaene biosynthesis | Private | (74) |
| **9340** | pksI | Bacillaene biosynthesis | Private | (74) |
| **9345** | pksJ | Bacillaene biosynthesis | Private | (74) |
| **9350** | pksL | Bacillaene biosynthesis | Private | (74) |
| **9355** | pksM | Bacillaene biosynthesis | Private | (74) |
| **9360** | pksN | Bacillaene biosynthesis | Private | (74) |
| **9365** | pksR | Bacillaene biosynthesis | Private | (74) |
| **9370** | pksS | Bacillaene biosynthesis | Private | (74) |

